# Sustained oscillations of epithelial cell sheets

**DOI:** 10.1101/492082

**Authors:** Grégoire Peyret, Romain Mueller, Joseph d’Alessandro, Simon Begnaud, Philippe Marcq, René-Marc Mège, Julia M Yeomans, Amin Doostmohammadi, Benoît Ladoux

**Author notes:** These authors contributed equally to this work (Grégoire Peyret, Romain Mueller, Joseph d’Alessandro).

## Abstract

Morphological changes during development, tissue repair, and disease largely rely on coordinated cell movements and are controlled by the tissue environment. Epithelial cell sheets are often subjected to large scale deformation during tissue formation. The active mechanical environment in which epithelial cells operate have the ability to promote collective oscillations, but how these cellular movements are generated and relate to collective migration remains unclear. Here, combining in vitro experiments and computational modelling we describe a novel mode of collective oscillations in confined epithelial tissues where the oscillatory motion is the dominant contribution to the cellular movements. We show that epithelial cells exhibit large-scale coherent oscillations when constrained within micro-patterns of varying shapes and sizes, and that their period and amplitude are set by the smallest confinement dimension. Using molecular perturbations, we then demonstrate that force transmission at cell-cell junctions and its coupling to cell polarity are pivotal for the generation of these collective movements. We find that the resulting tissue deformations are sufficient to trigger mechanotransduction within cells, potentially affecting a wide range of cellular processes.

---

The formation of multicellular patterns of moving cells in living tissues is a hallmark of many developmental and pathological processes including morphogenesis, tissue regeneration, and tumourigenesis^1,2,3^. The ability of the cells to coordinate their motion over large scales^2,4^ enables the rapid transmission of mechanical information across the tissue, for example during tissue invagination driven by acto-myosin contractions^5^, gastrulation^6^, the propagation of velocity waves far from the free edge of migrating monolayers^7,8^ or the transmission of contact-guidance signals away from localised topographical cues^9^. It is now well established that physical constraints play a key role in the regulation of collective motion^13,14,15^ both *in vitro* and *in vivo* ^16,17,18^. Such collective behaviours are often accompanied by the emergence of oscillatory mechanisms such as calcium waves^20^, acto-myosin contractile activity^21^, and cyclic activation of signaling pathways (e.g. Notch, Fgf, and Wnt)^22^. These oscillatory behaviours occur at various time scales and play a major role in tissue reshaping^19^.

Several studies have reported the emergence of collective oscillatory movements in confined cellular monolayers *in vitro*. Epithelial monolayers of Madin-Darby Canine Kidney (MDCK) cells in circular confinement have been shown to convert otherwise chaotic flow into one coherent rotation^23^. Careful analysis further revealed regular pulsations in the radial direction that are uncoupled from these large movements^24^. These pulsations were further investigated in^25^ where a coupling of the cellular polarisation to local gradients of contractility was suggested as a potential mechanism. Here, we describe a novel form of coherent oscillations in confined human keratinocytes (HaCaT) and enterocytes (Caco2), where contrary to the aforementioned phenomena the oscillatory motion is the dominant contribution to the cellular movements. Moreover, we are able to reproduce the previously reported observations using MDCK cells, asserting the different nature of the phenomenon investigated here. This new mode of collective movement relies on the coordination of cell velocities along a direction that varies in time, at the scale of the whole monolayer. These specific geometric properties have unanticipated consequences in anisotropic confinement and thus provide a potentially powerful mechanism for pattern formation. These large scale movements lead to alternating phases of tissue compression and extension which are sufficient to trigger a localised mechanosensitive signalling pathway as determined by the nucleo-cytoplasmic shuttling of the transcription factor, Yes-associated protein (YAP), which is an important regulator of tissue growth, regeneration and tumourigenesis ^26,29,30^. We demonstrate that the origin of this behaviour lies in force transmission at cell-cell junctions and we present a computational model coupling cell motility to intercellular forces directly which is able to account in detail for our experimental observations.

## Results

### Confined human keratinocytes exhibit coordinated oscillations

Human keratinocytes (HaCaT cells) were deposited on micro-contact-printed square areas of fibronectin of size 500 × 500 µm surrounded by non-adherent surface (see Methods). Cell displacement and velocity fields were analysed using time-lapse microscopy and particle-image velocimetry (PIV) (Fig. 1a). Cells were initially sparse but progressively expanded within the square through cell proliferation with a doubling time of about 24 h until the tissue occupied all the available space (confluence stage). The corresponding velocity magnitude averaged over the whole domain initially increased over time, then plateaued for almost 20 h, finally dropping to a low value (Fig. 1b) in a way reminiscent of a “freezing” effect due to crowding ^31,32^.

**Figure 1:**
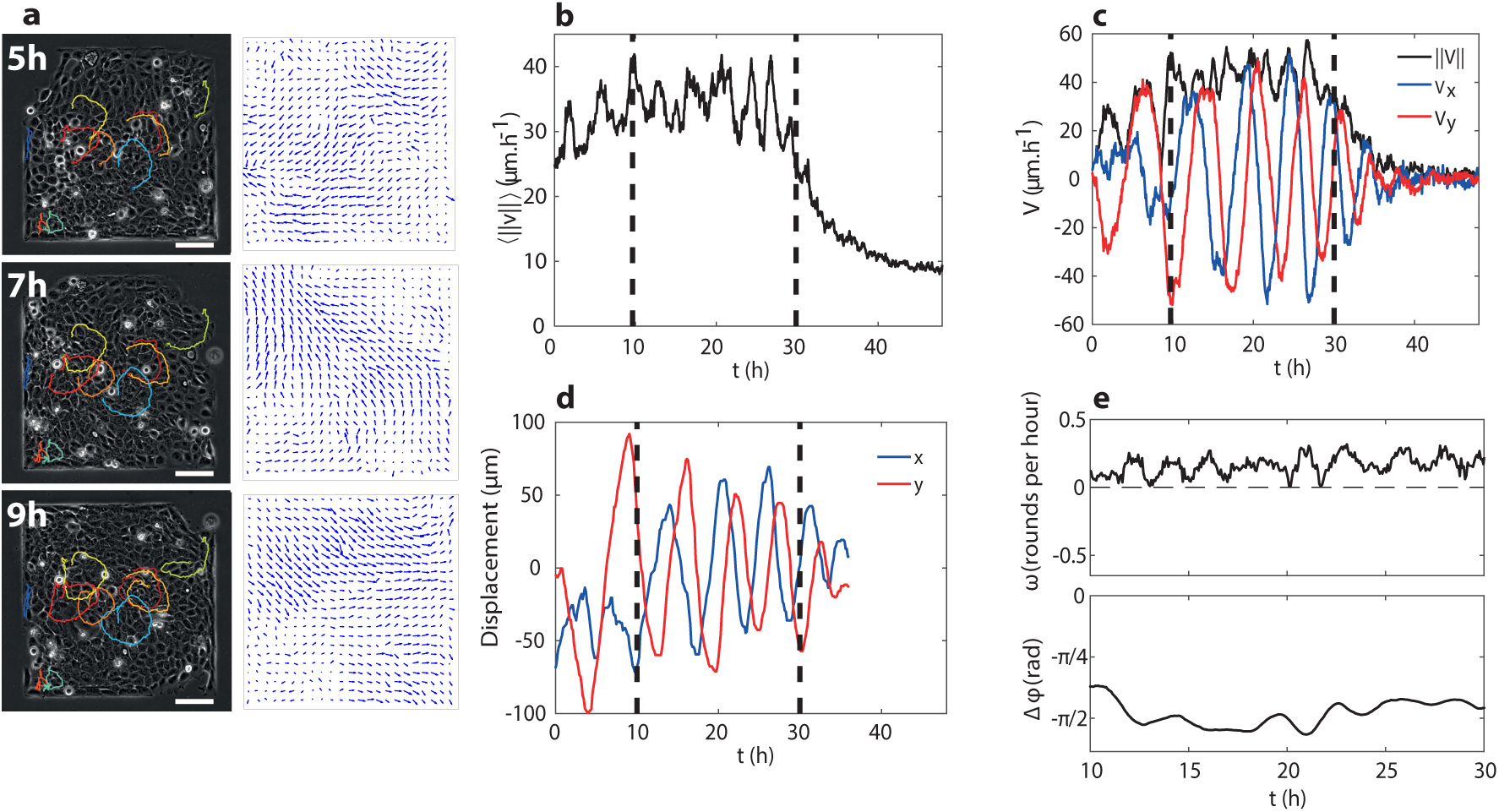
Coordinated oscillations of a confined epithelium. **a***, Left column: snapshots of a confluent HaCaT layer on a square pattern at various times and representative trajectories of single cells within the pattern (coloured lines)*. *Right: velocity fields from PIV measurement at the corresponding times*. *Scale bar is* 100 µm. **b***, Temporal evolution of the average velocity magnitude*. *The velocity increases slowly at first and then plateaus for almost* 20*h until it finally decreases*. *Vertical lines denote the period of time chosen to best highlight the oscillations*. **c***, Evolution of the two projected components — V_x_* = 〈*v_x_*〉_ROI_ *and V_y_* = 〈*v_y_*〉_ROI_ *— and the norm of the velocity* 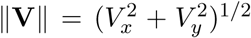. **d***, Displacement of a single cell in the centre of the domain*. **e***, Angular velocity of the average direction of the velocity (top) and phase-shift between V_x_ and V_y_ (bottom) within the time interval shown on figures b, c, and d*.

We then focused on the plateau stage where the cells were tightly packed but the movements were still large. Individual cells moved along parallel elliptical trajectories with little net relative motion of cells with respect to each other and no overall rotation of the tissue was observed (Fig. 1a,d and Suppl. Fig. 1). This led to the emergence of layer-scale coordinated movements in which all the cells moved together in a direction that rotated slowly with time (see Fig. 1a and Suppl. Video 1). As the cells moved collectively they pushed each other against the domain boundary, leading to a decrease of their two-dimensional effective area and to a local increase of the density. The oscillatory nature of these collective movements was best revealed by considering the total velocity 〈**v**〉_ROI_ averaged over a region of interest (ROI) of 50 × 50 µm at the centre of the pattern. While the magnitude ||〈**v**〉_ROI_|| was almost constant and non-zero, the individual components 〈*v_x_*〉_ROI_ and 〈*v_y_*〉_ROI_ showed clear oscillations (Fig. 1c) with a phase shift of π/2 (Fig. 1e bottom), indicating that 〈**v**〉_ROI_ rotated at constant magnitude inside the region of interest. The angular velocity of this rotation had a mean value of about 1 rad · h^−1^ which translates to an oscillation period of about 6 h. There was a slow increase of the frequency with time (Fig. 1e top), which is likely due to a slow increase of the cell density. At later times, these movements gradually disappear (Fig. 1c), although residual flows are still observed (see Fig. 1b). We finally checked that the observed behaviour also appeared in other epithelial cell types by repeating our experiments with Caco2 human intestinal cells and indeed observed marked oscillations in squares of 200 µm (Suppl. Fig. 2).

### The geometry of confinement controls the properties of the oscillations

We then studied how changing the geometry of confinement affected the properties of the oscillations by depositing HaCaT cells in squares of sizes 100 µm, 200 µm, 500 µm, and 1000 µm. We characterised these changes by measuring the mean amplitude and period of the oscillations in 〈*v_x_*〉_ROI_ and 〈*v_y_*〉_ROI_. The period was found to grow linearly with the domain size (Fig. 2b and Suppl. Fig. 3). Because the magnitude of 〈**v**〉_ROI_ is approximately constant across the centre of the domain, this suggests that the reorientation cues come from the confinement boundaries where the cells are most deformed (Suppl. Fig. 1). The amplitude of the oscillations was found to be constant on a broad range of domain sizes but was reduced in smaller domains (Fig. 2a and Suppl. Fig. 3). We hypothesised that this reduction appeared for confinement sizes much smaller than the intrinsic length scale of the collective flows spontaneously generated by the epithelial cells. This was confirmed by measuring the velocity-velocity correlation length in unconstrained monolayers, which we found to be 600 – 1000 µm (Suppl. Fig. 4). This suggests that there is a subtle interplay between the size of the confinement and the natural correlation length of the cells within a tissue: in order for the oscillations to appear, these two lengths must be of the same order for the confinement boundaries to provide restoring cues allowing the direction of motion to slowly rotate. When the confinement size is much larger than the correlation length, those cues cannot be transmitted over the entire tissue, preventing the emergence of monolayer-scale movement. Conversely, when the domain is too small, large-scale collective flows of cells are screened. This is further seen in experiments using Caco2 cells: with a measured correlation length of 70 µm, oscillations appear in 200 µm but not in 500 µm squares.

**Figure 2:**
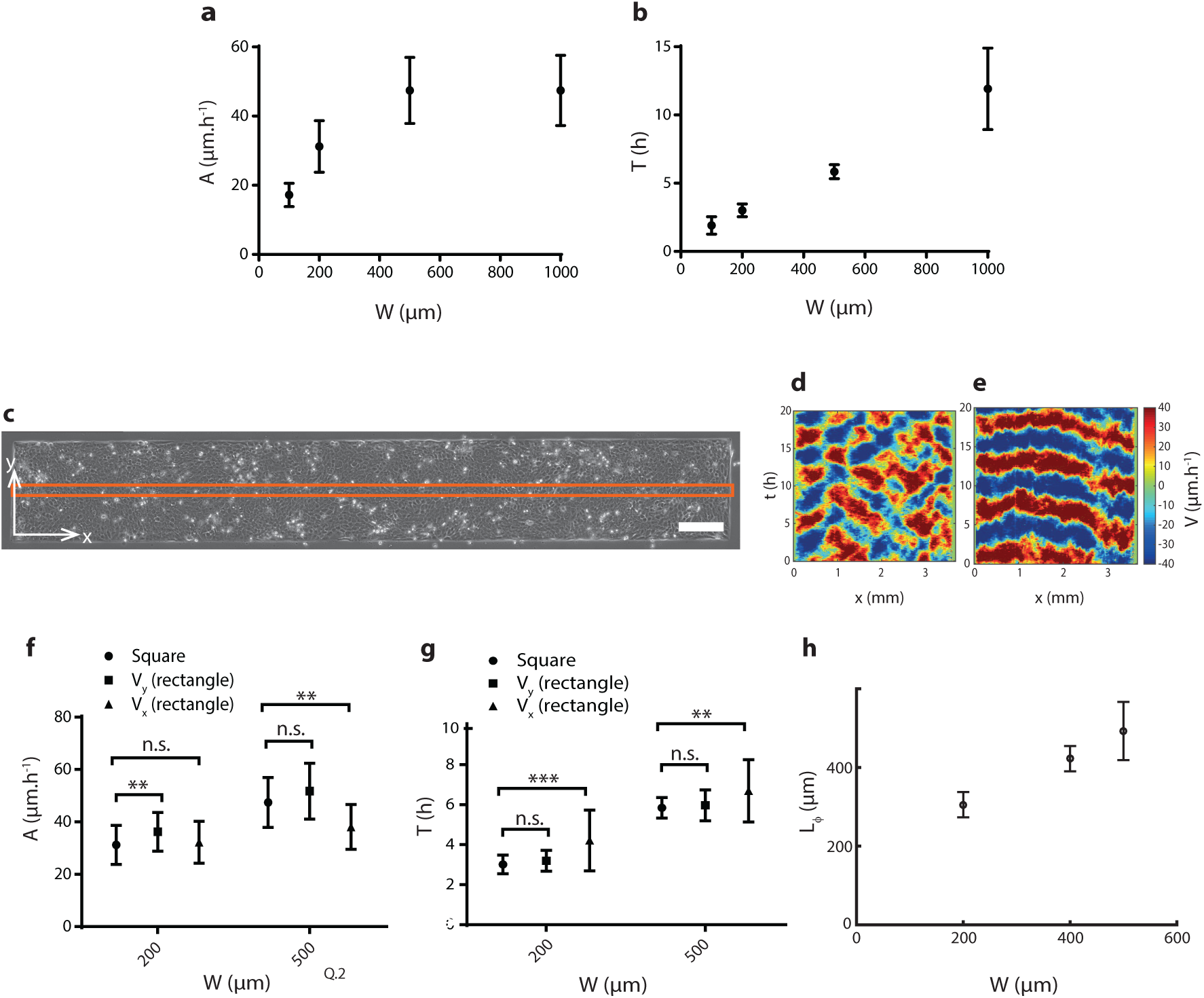
Geometry dependent coordinated oscillations. **a** *and* **b***, Amplitude A and period T of the oscillations as a function of the size of the square confinement*. **c***, Snapshot of rectangular confinement of size L* × *W* = 3500 × 500 µm. *The orange frame represents the region of interest used in figures d, e, and h*. *Scale bar is 200* µm. **d***, Spatiotemporal map of V_x_* = 〈*v_x_*〉_ROI_ *along the long axis shows patches of correlated motion with a coherence size of about* 500 µm. **e***, Spatio-temporal map of V_y_* = 〈*v_y_*〉_ROI_ *along the short axis*. **f** *and* **g***, Amplitude A and period T of the oscillations in squares and rectangles as a function of the smallest dimension of the confinements*. **h***, Phase coherence length L_ϕ_ as a function of the width of the rectangular pattern*. *L_ϕ_ is defined as the distance over which V_x_*(*x, t*) *reverses direction*. *All plots, mean (SD) from 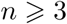 experiments*. *n*.*s*.*: not significant, *p<* 0.05*, **p<* 0.01*, ***p<* 0.001 *from a two-sample Kolmogorov-Smirnov test*.

To investigate this interplay further, we next considered anisotropic confinement and introduced high aspect ratio rectangular domains of width *W* = 200 µm, 400 µm, and 500 µm and length *L* =2, 000 − 3, 000 µm (Fig. 2c). Similar to square confinements, the cells exhibited large patches of collective motion. We then measured 〈*v_x_*〉_ROI_ and 〈*v_y_*〉_ROI_ over a rectangular region of interest (shown on Fig. 2c), averaging over the width of the rectangle. The velocity component in the short direction, 〈*v_y_*〉_ROI_, showed long-range coordination along the whole domain and oscillated between positive and negative values at each fixed position (Fig. 2e). In contrast, the velocity component along the long axis, 〈*v_x_*〉_ROI_, was not correlated over the whole domain but rather formed alternating patches of coordinated motion (Fig. 2d). Such velocity pattern corresponds to the cells moving in the rectangle in a fashion reminiscent of a standing wave in both directions with the wavelength equal to the width of the confinement *W* (Suppl. Video 2).

Most strikingly, the properties of this motion were set by the width *W* of the confinement while being apparently independent of the length: the period and amplitude of the oscillations closely matched those measured in squares of size *W* (Fig. 2g) and the typical size of the horizontal coordinated patches – as measured by the phase coherence length in Fig. 2h – matched closely with *W* as well. Even a migrating cell sheet with a free edge, hence confined in a single dimension, exhibits the same kind of behaviour, with clear coordinated oscillations in the direction of confinement and alternation of forward and backward moving zones in the perpendicular direction (Suppl. Fig. 2). This suggests that the period and amplitude of the oscillations are in fact independent of the exact shape of the confinement and only depend on the smallest confinement size. As a test of this hypothesis, we used circular confinement patterns instead of squares and recovered the oscillating behaviour with the same size-dependent properties (see Suppl. Fig. 2). This implies that the smallest geometric constraint acting on the system is able to impose a specific length scale to the tissue and selects associated patterns of motion. This pattern selection property might be of particular importance during development, which involves several shape formation and segmentation steps, and where collective oscillations have been shown to be crucial^33,34,35,36,37,38^.

### Epithelial oscillations lead to density fluctuations and mechanotransduction

Next we examined the impact of the oscillations on cell density fluctuations by measuring local densities. Imaging histone-GFP HaCaT cells (Fig. 3a) showed that the coordinated movements created large inhomogeneities in cell density distribution as shown by density maps (Fig. 3b). The relative density fluctuation magnitude Δ*ρ/ρ* = 18 ± 2% was consistent with the displacement-confinement relationship *Δρ/ρ* = 2Δ*w/w* ≃ 2Δ*x/W* = 2 × 8% (where *w* is the characteristic cell size and Δ*x* is the displacement amplitude, see Suppl. Fig. 1). To further assess the spatio-temporal fluctuations of cell densities, we computed the divergence of the velocity field from the PIV analysis (Fig. 3c). Positive divergence indicates local expansion of the tissue while negative divergence indicates local contraction. Interestingly, we observed that the oscillations were accompanied by local density variations and created alternating phases of compression and extension at any given point of the tissue.

**Figure 3:**
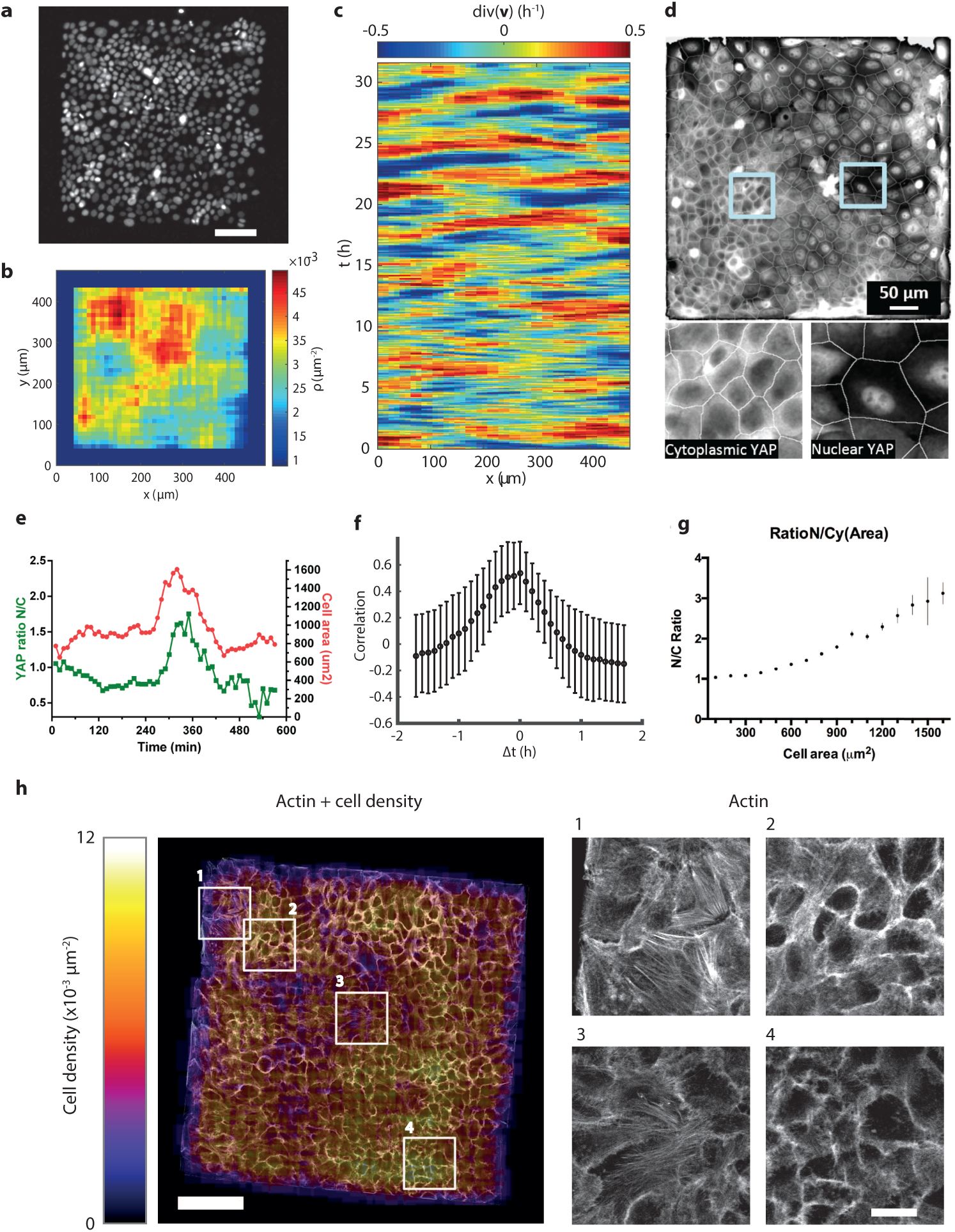
Oscillation-induced cell contractions lead to YAP translocation and actin reorganisation. **a***, Example cells expressing histone-GFP followed in time by live imaging*. **b***, Corresponding cell density map computed by counting the nuclei in a sliding window*. **c***, Spatio-temporal kymograph of the divergence of the velocity in a square of width W* = 500 µm*, showing alternating phases of compression and expansion*. **d***, Immunostaining of YAP on a fixed sample of confined HaCaT cells with zoomed areas (marked with a blue square) illustrating cytoplasmic (left) and nuclear (right) YAP localisation*. **e***, Temporal evolution of YAP nucleo-cytoplasmic ratio (green) and cell area (red) for an example cell*. *The event of spreading at t* ∼ 300 *min corresponds to a sudden increase in YAP N/C ratio*. **f** *Time cross-correlation of the single cell area and YAP N/C ratio, averaged over the cell population in three independent experiments*. *Mean (SEM) for n* = 15 *cells*. **g** *YAP N/C ratio as a function of the cell area in fixed samples of confined epithelia*. **h** *Confocal picture of actin staining in the basal plane of fixed HaCaT cells confined on a square of width W* = 500 µm*, overlaid with the local cell density*. *Scale bar* 100 µm. *Right: zoomed areas (marked with a red and pale blue square) showing di*ff*erent organisations of actin fibres with respect to the local density (1,3, low density; 2,4, high density)*. *Scale bar* 20 µm. *Four pictures, same scale and contrast*.

To investigate the potential mechanotransductive effects of such fluctuations on biomechanical signals, we analysed the intracellular distribution of the transcription factor YAP, which is known to respond to compression ^29,45^. Various mechanical factors have been shown to convert phosphorylated YAP (retained in the cytoplasm where it is inactive) into its non-phosphorylated form (transcriptionally active and localised in the nucleus)^26,29,48^. We transiently transfected YAP-GFP into HaCaT cells and then deposited them on 500 µm square patterns in a 1:10 mixture with non-transfected cells and observed a high activation of YAP in regions of low densities whereas YAP appeared inactive in dense regions (Fig. 3d bottom). We quantified the YAP nucleo-cytoplasmic ratio (YAP-NCR) with respect to cell area of individual cells and observed significant fluctuations of cell area, which were followed closely by variations in the YAP-NCR (Fig. 3e and Suppl. Video 3). The temporal cross-correlation of both signals showed a maximum at Δ*t* = 0 (Fig. 3f), clearly indicating the direct relationship between cell area and YAP nuclear localisation in our system. We could not determine a delay between the two signals, which could be due to the very fast, mechanically-induced entry of YAP into the nucleus upon cell deformation^49^. We then considered wild-type cells in the oscillatory phase and imaged YAP by immunostaining. This allowed us to observe the large inhomogeneities in cell density created by the oscillations at a given time (Fig. 3d top) and to confirm the strong correlation between cell spreading and YAP nuclear translocation (Fig. 3g). Taken together, these results show that alternating phases of extension and contraction resulting from epithelial oscillations induce a strong and spatially resolved regulation of YAP localisation. Conversely, the inhibition of YAP transcriptionnal activity using verteporfin did not affect the oscillations (Supp. Fig. 5). It suggests that YAP translocation is only a molecular readout of the oscillations but might not retroact on them.

Those changes in local cell density were also accompanied by out-of-plane deformations: squeezed cells in high density areas were taller, while at low density the cells were flatter (Supp. Fig. 6). Finally, those variations of density and cell shape were also correlated to changes in the cytoskeleton organisation: in the low density areas, cells formed parallel, cell-scale, actin fibres suggesting high tensional state, while in more compressed zones, F-actin was restrained to the cell cortex. Hence the mechanical state and active force production of the tissue were sensitive to the fluctuations in cell density, a clue of a potential feedback from and on cellular movements, which is necessary for the maintenance of the oscillations.

### Local cell-cell forces, motion and polarity are coupled to the collective dynamics

To gain further insights into the dynamics of the oscillations we then used traction force microscopy^39,40^ to measure the forces exerted on the substrate by the cells. In general the traction forces were heterogeneous within the tissue and their largest values were predominantly located at the edges of the pattern and directed inwards (Fig. 4a). Intermittent patches of high force were observed, as a result of lamellipodia transiently pulling on the substrate. Combining traction force microscopy with Bayesian inference stress microscopy^46^, we next measured the pressure *P* = −Tr(*σ*)*/*2 within the tissue and found it to be under homogeneous tension, except for a thin edge layer where the tension drops to zero (Fig. 4b). Actin staining further showed that the tissue tension is supported by the actin cytoskeleton and transmitted across cell-cell junctions through small actin digitations (Fig. 4c). As opposed to other types of cell-cell contacts in flat epithelia that are associated with parallel oriented F-actin bundles, radially-oriented F-actin bundles are reminiscent of high tension at cell-cell contacts as previously described^47^.

**Figure 4:**
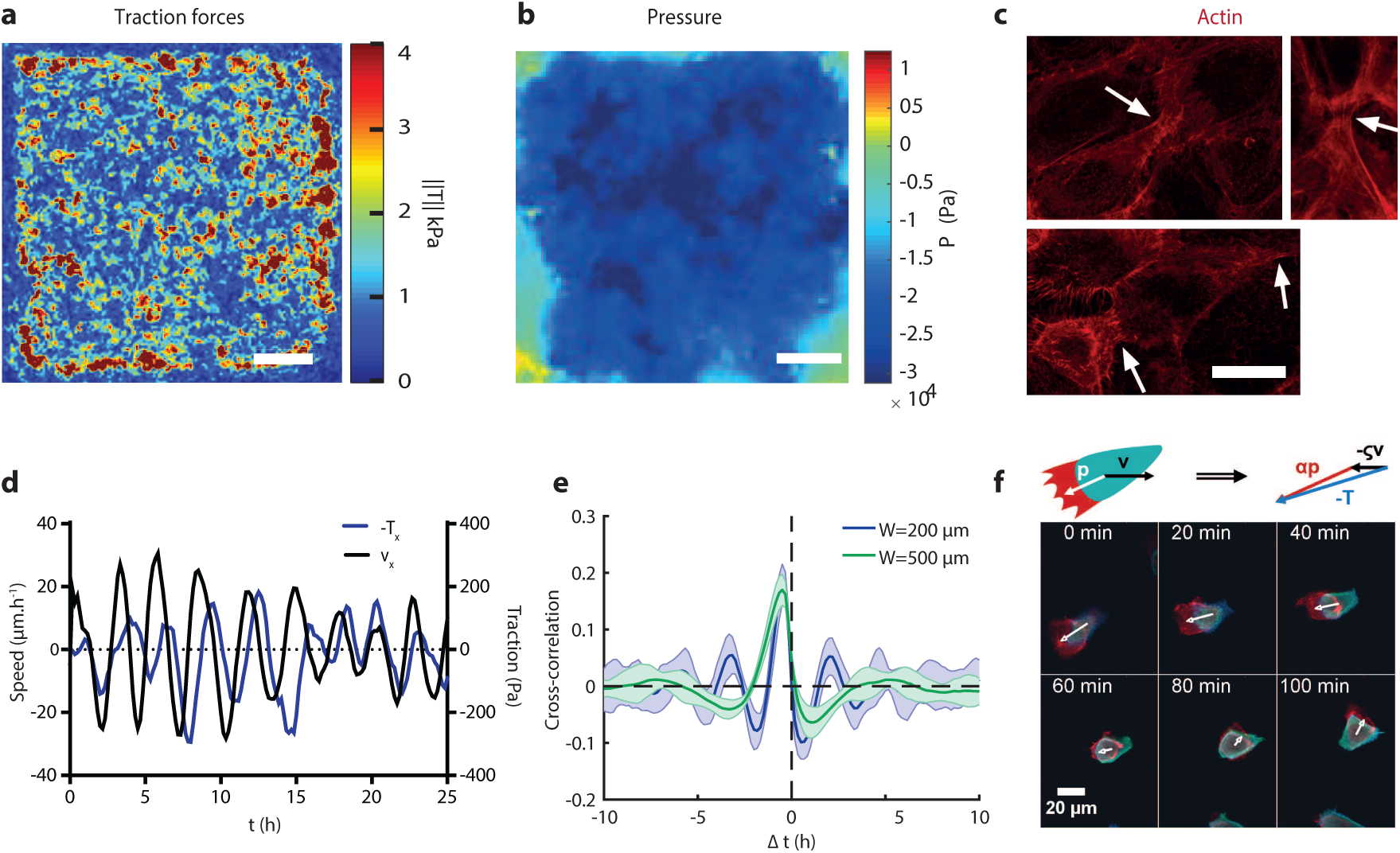
Traction force and cell polarity are coupled to cell motion. **a***, Map of the magnitude of the traction forces*. *Scale bar* 100 µm. **b***, Map of the pressure P* = −Tr(*σ*)*/*2 *within the confined tissue*. *Scale bar* 100 µm. **c***, Confocal images of actin staining in fixed HaCaT cells fixed during the oscillation phase*. *The arrows denote cell-cell junctions under tension*. *Scale bar* 20 µm *(same scale for three pictures)*. **d***, The averages of v_x_ and* −*T_x_ in a central square ROI, projected on the x-axis, oscillate with similar period but are out of phase*. **e***, Normalised cross-correlation between* **v** *and* −**T** *as a function of the time-lag Δt*. *The correlation has a marked peak at Δt* =0.5 *h for both W* = 200 *and* 500 µm. *Mean (SD) of n* = 11 *and* 8 *respectively*. **f***, Top: Schematic of the cell polarity: the orientation of the lamellipodium (red) with respect to the cell body (turquoise) defines the polarity (vector* **p***), which can be in a di*ff*erent direction from the velocity* **v**. *The traction force* **T** *is a linear combination of the two*. *Bottom: Example of a reorientation event (fluorescent cell embedded in a non-fluorescent confluent mono-layer)*. *F-actin in the basal plane (z* =0*, red) and top of the cell (z* = +7 *and* +8 µm*, green and blue), visualised by confocal microscopy*. *The spatially shifted basal actin shows the protrusion orientation, used to define the cell polarity and denoted by a white arrow*. *The polarity, initially directed towards the bottom-left corner, realigns in the direction of motion (towards the upper-right corner) over time*. *Scale bar* 20 µm.

It was recently argued^25^ that the traction force and the velocity are not necessarily aligned. Indeed we observed that the angle between these two directions followed a flat distribution when averaged over the whole experimental time (Suppl. Fig. 9). However, at fixed times and averaging only over a centred region of interest, we found that the traction forces and the velocity are correlated with a well-defined delay (Fig. 4d). The cross-correlation of the two signals showed a clear peak at Δ*t* = −0.5 ± 0.1 h both for 200 µm and 500 µm squares (Fig. 4e). This suggests that cells adapt their motion to the surrounding tissue by polarising in the direction where they are pulled by their neighbours, with a typical realignment time of the order of 30 min. Therefore, in contrast to the conclusions presented in^25^, we find that the traction forces and the local velocity were not simply uncorrelated but rather showed alternating phases where the traction force was aligned to the motion of the cells and phases where it was pointing in the opposite direction.

We could visualise this phenomenon directly by mixing wild-type cells with a small proportion of HaCaT expressing lifeact-GFP and imaging them using a confocal spinning disk microscope. This allowed us to see clearly the lamellipodia created at the basal side of the marked cells and to observe several reorientation events in which a cell was pulled by the surrounding tissue in the direction opposite to its lamellipodium and then reoriented its polarity so as to align it with its direction of motion (Fig. 4f).

### Contributions of cellular-level mechanisms to the oscillations

Next we asked which mechanisms at the cell level are able to modulate the oscillations. We began by studying the role of cell motility through drug inhibition of cytoskeletal activities. To this end, actomyosin contractility was targeted by the inhibition of myosin-II processivity using blebbistatin, while protrusion formation was suppressed by the inhibition of Arp2/3-mediated branched actin nucleation using CK-666.

Upon addition of blebbistatin at high concentration (50 µM), the cell speed fell rapidly to a very low value (Suppl. Fig. 7). The oscillations, however, survived albeit with larger period and smaller amplitude (Fig. 5a-b). The approximately constant product of *period* × *amplitude* ≈ 80 µm suggests that only the migration speed was affected by the drug. This implies that, contrary to what was reported for the “breathing” oscillations^25^, actomyosin contractility is not directly responsible for the observed oscillations even though it is necessary to drive the system out of equilibrium. Upon addition of CK-666 (100 µM), the speed first slightly dropped and then gradually decayed over more than 30 h (Suppl. Fig. 7). While none of the oscillations’ properties was affected immediately after the drug addition, the long term slow down was accompanied by a decrease of the total velocity and a fading out of the oscillatory behaviour (Suppl. Fig. 7). This is not surprising because the protrusive activity is necessary for the cells to direct their motion so that its suppression is expected to result in the loss of the directed collective motion of the cells. However the decay of the oscillations is slow suggesting that protrusive activity is not directly responsible for the emergence of oscillations. Together, these findings show that while actomyosin contractility and protrusion formation are required for fueling cell movements they do not directly lead to the emergence of oscillations.

**Figure 5:**
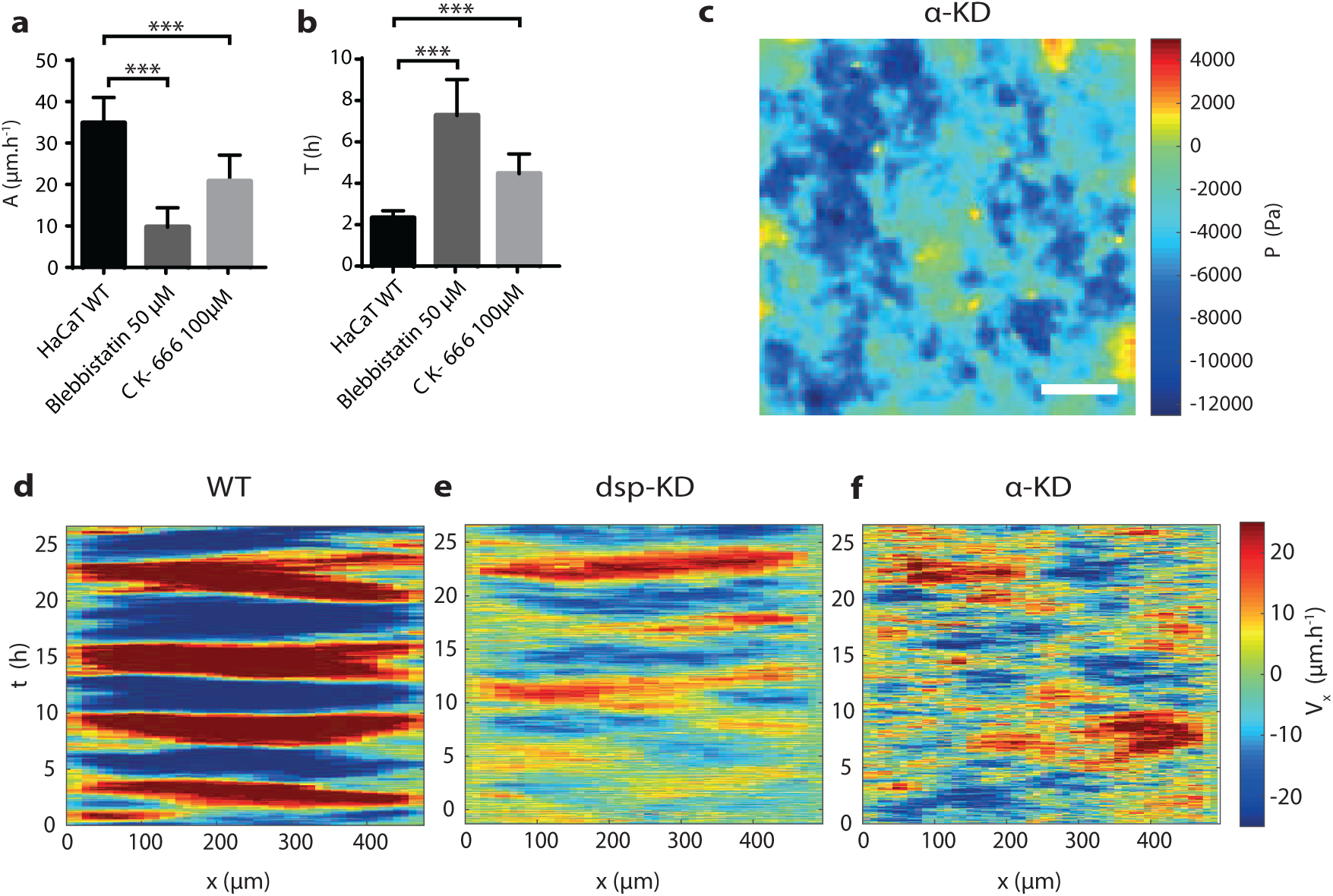
Genetic and molecular perturbation experiments. **a** *and* **b***, Amplitude A and period T of the oscillations in* 500 µm*-sized squares, for control cells and with blebbistatin at* 50 µM *and CK-666 at* 100 µM. *Mean (SD) from n* =5 *patterns*. **c***, Pressure map in α*−*catenin knocked-down cells*. *Scale bar* 100 µm. **d–f***, Kymographs of V_x_* = (*v_x_*}_ROI_ *for wild-type (d), desmoplakin knocked-down (e) and α*−*catenin knocked-down (f) cells*. *n*.*s*.*: not significant, *p<* 0.05*, **p<* 0.01*, ***p<* 0.001 *from a two-sample Kolmogorov-Smirnov test*.

We then tested the contribution of cell-cell junctions to the oscillations by disturbing them using gene silencing via stably expressed shRNAs, targeting either desmoplakin or *α*-catenin. Desmoplakin is a critical component of desmosomes, which are cell-cell adhesion structures linked to the intermediate filament cytoskeleton^41,42^, while *α*-catenin is involved in the cadherin-actin linkage at the adherens junctions, where it is known to stabilise the junctions and participate in mechano-chemical signal transduction^43,44^. Desmoplakin-knocked-down HaCaT cells (dsp-KD HaCaT) displayed oscillatory motion with period similar to that of control cells but the velocity field showed more marked fluctuations and the amplitude of the movement was slightly decreased (Fig. 5d-e). More strikingly, *α*-catenin-knocked-down HaCaT cells (*α*-KD HaCaT) exhibited completely disordered movements: cells were able to slip past each other as shown by the low value of the correlation length – ~ 30 µm, which resulted in random trajectories and no measurable oscillation (Fig. 5f and Supplementary Video 4). This indicates that force transmission at adherens junctions is essential for the oscillations to emerge.

Disrupting the force transmission at the cell-cell junctions also impacted the internal stress *σ* of the tissue. In the wild-type cells, the pressure *P* = −Tr(*σ*)*/*2 is negative and homogeneous except in a thin edge layer (Fig. 4b) showing that the tissue is always under tension. In *α*-KD HaCaT cells, *P* is still negative but 3 to 4 times lower on average, and its spatial distribution showed marked inhomogeneities (Fig. 5c). This indicates that force transmission is strongly affected and supports the idea that the force transmission across cell-cell junctions is a critical factor in the appearance of oscillations.

### A simple coupling between the interface forces and the cell polarity reproduces the experimental results in a generic model

We now show that a computational model that couples mechanical forces at the cell-cell interfaces to the individual cell polarity is capable of reproducing our experimental results. We describe individual cells as deformable and compressible active particles using a phase-field approach^50,51^ that models the interface of the individual cells explicitly and incorporates cell compressibility and inter-cellular forces directly (see Methods). Besides its deformable interface, each cell *i* has a velocity **v***_i_* and is associated with a vector **p***_i_* describing its polarity and defining the direction of the propulsive thrust it exerts on the substrate^52^. We consider over-damped dynamics and write the force balance condition 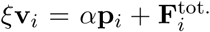, where *ξ* is a friction coefficient, *α* is the strength of the motility, and 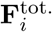. is the total force acting on one cell’s boundary which has contributions from steric repulsion and friction with both neighbouring cells and the walls, see Methods.

We now specify the dynamics of the polarity following our experimental observation that the cell velocity is correlated with the traction force with a well-defined delay. For simplicity we fix **p***_i_* = (cos *θ_i_*, sin *θ_i_*) and define the following diffuse dynamics for the angle

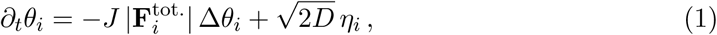

where Δ*θ_i_* ∈ [−*π, π*] is the angle between **p***_i_* and 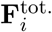, and *η_i_* is a Gaussian white noise. The positive constants *J* and *D* are the strength of the alignment torque and the rotational diffusivity, respectively. Equation (1) describes the diffusive alignment of the polarity of the cells to the total force on their interfaces and we show below that it induces the correct correlation of the velocity with the traction force. This formulation differs strongly from the model of Notbohm *et al*.^25^ which crucially relies on prescribing an additional reaction-advection equation with a predefined relaxation time for an unknown chemical (such as phosphorylated myosins) controlling the strength of the isotropic active stress, resulting in an indirect coupling of the polarity direction to changes of local cell area. In contrast, our model does not require the introduction of an internal chemical and couples the polarity directly to the resultant pushing/pulling and tangential – frictional – forces exerted on the surface of the cell.

Simulations of confluent monolayers in square confinement show the emergence of sustained oscillations (Fig. 6a,b and Suppl. Video 5). As the cells move towards one of the edges they start to compress each other resulting in an increase of the steric repulsion force in the direction opposite to their motion, which then induces the reorientation of their polarity. We can pinpoint the crucial importance of the alignment mechanism in the generation of the oscillations by considering changes in the period of the oscillations as the coupling *J* is varied: for fixed box size, the period *T* scales as *J*^−1^ and diverges as *J* vanishes, showing that the oscillations effectively disappear when the alignment mechanism is absent (Fig. 6d). This reflects our *α*-catenin knock-down experiments, where we found that force-transmission through cell-cell junctions is crucial to the tissue oscillations Moreover, we finally check that our dynamics for the polarity induces the correct alignment between the velocity and the traction force. Indeed, the cross-correlation between **v***_i_* and −**T***_i_* shows a clear peak located at a non-zero time difference and whose location is independent of the box size *W*, see Fig. 6f. This indicates that, similarly to our experimental observations, the velocity follows the traction force with a well defined delay in the simulations.

**Figure 6:**
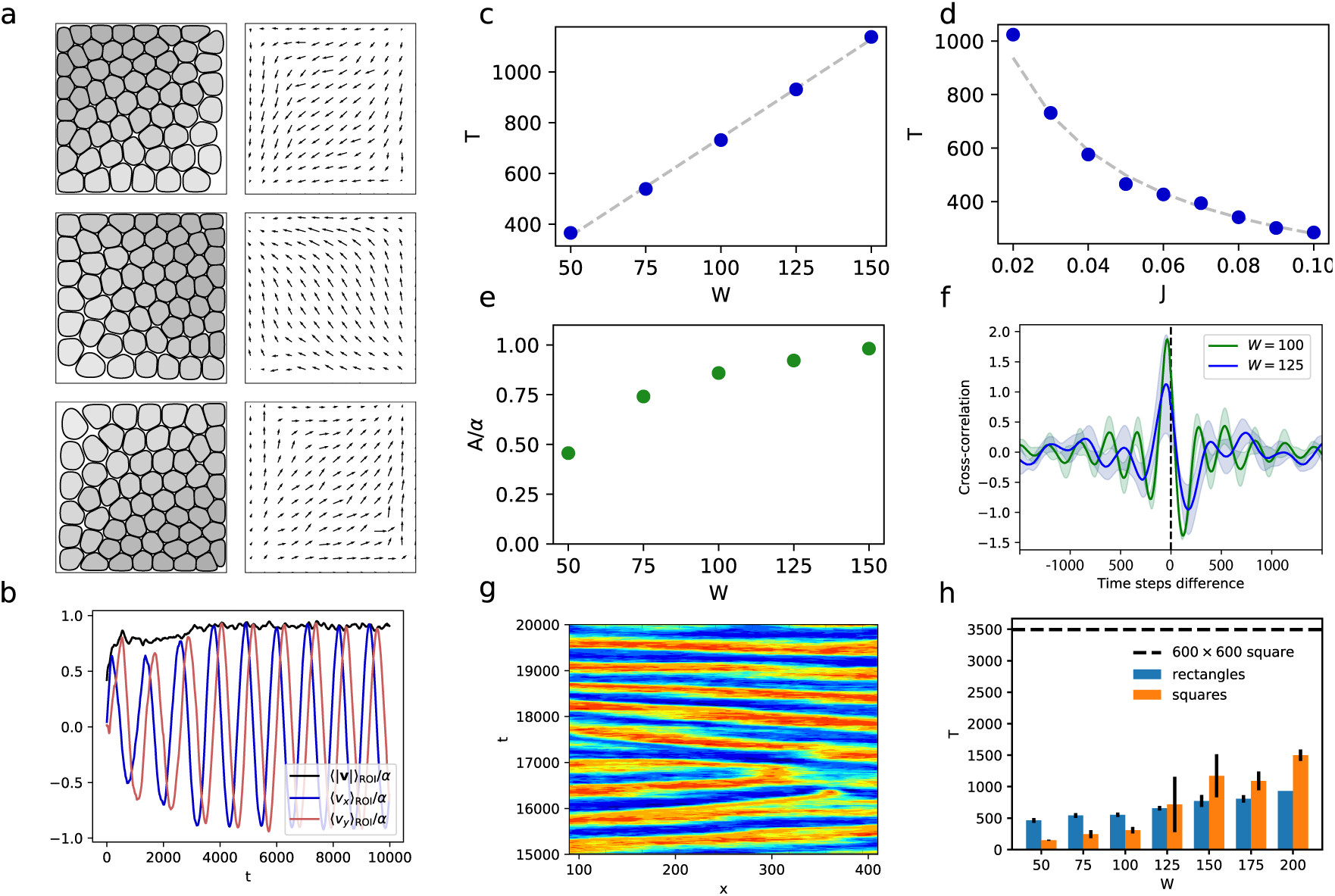
Computational model of collective cell motion. **a***, Snapshots of simulations showing an oscillating tissue at different times*. *The left-hand figures show the individual cells, where darker cells are more compressed, while the right-hand figures shows the corresponding velocity field*. **b***, Velocity projected on the x and y axes for the same system*. **c** *and* **d***, Dependence of the period T on the system sizes W and alignment parameter J, see equation* (1). *The grey dashed lines are least-square fit of T* ∝ *W and T* ∝ *J*^−1^. **e***, Dependence of the amplitude over active speed A/α for different system sizes W*. *The grey dashed line is a least-square fit of T* ∝ *W*. **f***, Cross-correlation between* **v** *and* −**T** *for two system sizes*. *Mean (SD) from n* =5 *simulations*. **g***, Kymograph of* 〈*v_x_*〉_ROI_ *averaged over the short direction for a rectangular box of size* 600 × 100 *with non-zero friction at the walls*. **h***, Period of the standing waves in rectangles compared to the period of oscillation in squares of size identical to the small dimension of the rectangle*. *The period of oscillation in a square of size* 600 × 600 *corresponding to large dimension of the rectangles is shown for comparison*. *Mean (SD) from 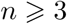 simulations*.

In agreement with the experimental results, the period of oscillations grows linearly with the system size while the amplitude grows at first and then reaches a plateau (Fig. 6e). The linear dependence of the period occurs because the speed of propagation of the velocity oscillations does not depend on the system size. Moreover, the amplitude is reduced in boxes smaller than the correlation length because the migration speed is decreased by the proximity of the walls, which hinders the motion of the cells. The amplitude of displacement also depends linearly on the system size (Suppl. Fig. 10d). Similarly, simulations of cells in rectangular confinements reproduce the experimental observation of standing waves along the short dimension of the channel (Fig. 6g). The period of these oscillations is dictated by the smallest dimensions as in the experiments and corresponds to the period obtained in squares of similar dimension (Fig. 6h). In conclusion, our model provides additional evidence that the interplay between force transmission at cell-cell interfaces and the reorientation of the cells lies at the origin of the oscillations.

## Discussion

In this paper, we have evidenced a novel mode of collective motion for epithelial cells under confinement. In isotropic geometries, oscillations that are coordinated over the whole confining domain emerge spontaneously. A striking feature of this phenomenon is that it does not rely on the phase synchronisation of individual oscillators – as in most biological collective oscillations – but is an emergent property of the group. We identified a coupling of the cell polarity to the forces transmitted through adherens junctions as a key ingredient. Determining the molecular mechanisms underlying this coupling is an open challenge for future research: the polarity could reorient following signals induced by forces exerted on the cell-cell junctions, in a manner reminiscent of contact inhibition of locomotion^56^; By contrast to previous studies^7,12^, we show that large-scale force transmission through cell-cell junctions is coupled to local polarity mechanisms at the single cell level to produce efficient coordinated movements. In large unconstrained tissues, these active mechanisms lead to collective patterns of motion, driven by the coordination of velocities and feedback from resulting large cell density fluctuations. Such spontaneous large-scale motions are critical during wound healing and epithelial gap closure^3,57,58^. We indeed show that the emergence of oscillatory cell motions can favour the closure of non-adherent epithelial gaps in vitro (See Supplementary Video 6). Such observation could shed new light on the mechanisms at play during in vivo wound healing. The collective movements come with a broad distribution of time and length scales (Supplementary Figure 4). Thus, the confinement here simply enhances a single wavelength among those. Interestingly, the different behaviours exhibited by various epithelial cell types may reflect more general rheological properties of the monolayers themselves. For instance, HaCaT – skin – cells exhibit coordinated movements on the millimeter scale, while this length scale is of the order of 100 µm for Caco2 intestinal cells. A noticeable aspect of the phenomenon is that due to the rotating character of the polarity axis, the two directions of movements are coupled. Hence, confinement in one single direction sets the length scale of the oscillations in the other direction, even in the presence of a freely moving edge with leader cells (Suppl. Fig. 2). As a result, a highly anisotropic tissue exhibits very regular patterns of velocities along the long axis, whose scale is determined by the short axis. One could thus imagine such a mechanism to be relevant in segmentation processes of high aspect ratios, such as the somite formation^22,59^.

Our molecular perturbation analyses clearly show the crucial role of adherens junctions in modulating the collective oscillations. They participate in the coordination of the cell polarities and also provide a route for the effective transmission of mechanical stresses to generate collective oscillations across the tissue. The observation of YAP translocation accompanying the oscillation-induced cell deformations further shows that the mechanical signals are sufficient to transduce mechano-chemical responses. Indeed, the deformations induced by these coordinated epithelial oscillations might provide a clock for the activation of cellular processes, or define their location. In this regard, it has been shown previously that patterns of cell differentiation can follow patterns of mechanical signals^60^. Beyond that, we anticipate that if such oscillations are able to emerge on sufficiently soft surfaces, they could also promote mechanical movements and tissue invagination in the third dimension, an important but poorly understood process during embryonic development.

## Methods

### Cell culture and reagents

HaCaT cells, MDCK cells and CaCo2 cells were cultured in DMEM supplemented with 10% foetal bovine serum and 1% penicillin-streptomycin in incubator at 37°C with 5% CO_2_ and humidification. Protein expression in HaCaT cells was depleted by retrovirus-mediated introduction of shRNA into cells, as described previously ^57^. HaCaT cells stably expressing Life-Actin-GFP and histone2B-GFP were established by transfection of pCAG-mGFP-Actin plasmid^62^ (gift from Ryohei Yasuda, Addgene plasmid # 21948) and pRRlsinPGK-H2BGFP-WPRE (gift from Beverly Torok-Storb, Addgene plasmid # 91788) respectively. For visualisation of human YAP1 in live cells, wild-type HaCaT cells were transiently transfected with the pEGFP-C3-hYAP1 plasmid^63^ (gift from M. Sudol, Addgene plasmid # 17843). For immunofluorescence, wild-type HaCaT cells were fixed with formaldehyde 4%, then labelled with an anti-human-YAP1 antibody (sc-101199, Santa Cruz Biotechnologies) or phalloidin-Alexa 568 (Life Technologies, Paisley, UK). For inhibition of contractility and actin polymerization experiments, drugs (blebbistatin 50 µM and CK-666 100 µM respectively) were added at the beginning of the experiment, approximatively 12–18 hours after seeding the cells on the patterns.

### Micro contact printing

Functionalisation of substrates was done using micro contact printing techniques. Master silicon moulds containing the patterns were made using SU-8 photoresist resin and were exposed for two hours to silane gas before being used. Stamps for patterning were made using soft lithography techniques. PDMS (Sylgard 184, Dow Corning) was mixed with current agent at a ratio of 1:10, degassed and poured over the wafer. PDMS was cured for two hours at 80^°^C and removed from the mould before being cut. Plastic Petri dishes and glass bottom Petri dishes were first coated with a thin layer of PDMS before printing. A drop of 1:10 mixed and degassed PDMS was put on the center of the substrate. A spin-coater with a first cycle of 18 s at 500 rpm and a second cycle of 1 min at 5000 rpm was used to spread the PDMS over the substrate. The printing of the fibronectin was performed as previously described ^58^. Briefly, PDMS stamps were incubated for 45 min with a 50µg/ml fibronectin solution in which a part was conjugated with a Cy3 or Cy5 dye to control the quality of the transfer. The substrates were isolated for 10 min in a UV-ozone machine before printing. Stamps were rapidly soaked 1 s in distilled water and completely air dried before being applied for 1 min on the substrate. PDMS stamps can be sonicated in ethanol and washed in MQ water before being reused. The non-printed surface of the substrate was passivated with a 2% pluronics F-127 solution for 2h. After being washed several times with PBS, cells were seeded on the substrate at high concentration and washed after 10–15 min. The substrate was placed in the incubator overnight to let the cells spread and imaged the next morning. The patterns were chosen such that the cells were almost confluent at the beginning of the experiments.

### Traction force microscopy

The substrates were prepared as described in a previous reference^57^. Soft silicon substrates were made by mixing CyA and CyB at a ratio of 1:1 and directly poured on glass bottom Petri dishes (fluorodish) in order to obtain a 100 µm thick layer. The substrate was cured at room temperature overnight on a flat surface. The surface was silanized using a solution of APTES at 5% in absolute ethanol for 5 min. The substrate was then washed with absolute ethanol and dried at 80^°^C for 10 min. 200 nm carboxylated fluorescent beads (Invitrogen) were added in a deionized water solution at 1:500 for 5 min, washed with deionized water and dried at 80^°^C for 10 min. For these substrates, we used a modified micro contact printing technique ^17^. The PDMS stamps were incubated with a fibronectin solution the same way as for the patterning on the PDMS. After being dried, the stamps were used to print a polyvinyl alcohol membrane (PVA). The printed piece of membrane was then deposed on the soft substrate for 30 min and then dissolved with a 2% F-127 pluronics for 2 h. The cells were seeded at high concentration (1 million per Petri dish) for 20–30 min and washed with PBS when the desired concentration was reached. At the end of the experiment, cells were removed with 500 µL of 10% SDS in the media. A stress free image of the substrate were taken to be compared to the other images.

### Live cell imaging and analysis

Live imaging was performed with a 10X objective on a BioStation IM-Q (Nikon) at 37^°°^C and 5% CO2 with humidification. One image was taken every 4 min for the experiments which did not need fluorescence. For the TFM experiments, a phase-contrast image and a *z*-stack of the fluorescent beads were taken every 10 min. Images were first processed with ImageJ to obtain the best focus plane for each time point (Stack Focuser plugin), stabilized (Image Stabilizer plugin), and background beads were removed. To analyze cellular movements, velocity fields were calculated by PIV analysis with MATPIV 1.6.1, a Matlab (the Mathwork) implemented script. An interrogation window of 32 (20.7 µm) or 64 pixels (41.4 µm) was selected for 4 min or 10 min per frame respectively, with an overlap of 50 to 75%. Vectors higher than a speed threshold manually determined were removed, and a local median filter was applied. Gradient of the velocity field was calculated with a method adapted from^61^. Briefly, based on the Green-Ostrogradsky theorem, we integrated the velocity vectors on a ring of radius 37.5 µm. This method has the advantage of reflecting the movement at the tissue scale more than local fluctuations, and is less noise sensitive. We calculated the strain-rate tensor from the gradient of the velocity field as 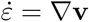. Correlation length was calculated as previously described^65^. Briefly, at a given time *t*, the (two-dimensional) spatial velocity autocorrelation function was computed in the Fourier space, then averaged azimuthally. The correlation length *ξ*(*t*) was obtained from a fit with a stretched exponential function *C*(Δ*r*) = exp(− (Δ*r/ξ*)*^ν^*). In the text we quote the range of typical values taken by this function in an unconstrained monolayer (HaCaT cells) or in a 500 µm square (Caco2 cells). For TFM experiments, the substrate displacements also were calculated with MATPIV 1.6.1. Interrogation windows were set to 24 pixels (15.5 µm) and 75% overlap. A median filter was applied to the displacement vector field. Traction force fields were computed with a FTTC plugin on ImageJ with a regulation factor set to 1 × 10^−10^. The stress within the tissue was calculated using Bayesian Inversion Stress Microscopy^46^ with a regularisation parameter Λ = 10^−6^.

### Statistics

Differences between data were assessed using two-sample Kolmogorov-Smirnov tests implemented in Matlab. On the plots, n.s.: not significant, \**p<* 0.05, \*\**p<* 0.01, \*\*\**p<* 0.001.

### Computational model

We consider a two-dimensional tissue and describe each cell *i* independently by a phase-field *ϕ_i_*, where *ϕ_i_* ≃ 1 indicates the interior of the cell and *ϕ_i_* ≃ 0 its exterior. Each cell is described implicitly and its boundary is defined to lie at the midpoint *ϕ_i_* =1*/*2. The phase-fields satisfy an overdamped dynamics given by

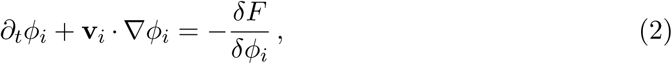

where 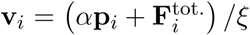 is the speed of the cell *i*, *F* is the free energy given below, *α* is the strength of the motility, *ξ* is a friction coefficient, and 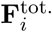 is the total force acting on one cell’s surface (including steric and viscous interactions, see below). The sole source of activity in the model is provided by *α* which induce a propulsion in the direction of the polarity **p***_i_*. Note that *ϕ_i_* is not conserved because we require the cells to be compressible.

The role of the total free energy *F* is both to maintain the cell integrity as well as to define interactions between cells. It can be decomposed as follows: a Cahn-Hilliard free energy term responsible for the stabilisation of the diffuse interfaces, a quadratic soft constraint enforcing area conservation, and finally two terms giving rise to repulsion forces between cells and with the confining walls. Following^51^, these contributions are defined as

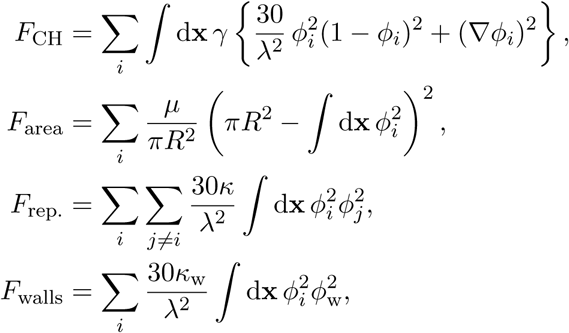

and we refer the reader to this reference for details. Note that because we only consider confluent tissues we do not introduce an adhesive force between cells. This is justified because an imbalance of repulsion forces has the same effect as an adhesion force in the force balance condition. Boundary conditions are implemented through the contribution *F*_walls_ as a repulsion between a fixed phase-field *ϕ*_w_ and the phase-fields *ϕ_i_*. The different parameters of the model are the following: cell stiffness (*γ* and *ξ*), compressibility (*µ*), repulsion strength (*κ* and *κ*_w_), polarity dynamics (*J* and *D*) and activity (*α*). Note that the interface width *λ* sets the basic length scale and is hence not considered as a parameter.

### Total force at the cell-cell interface

We write the total force experienced by a cell as 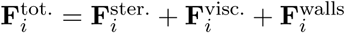, where the steric, viscous, and wall forces are defined as

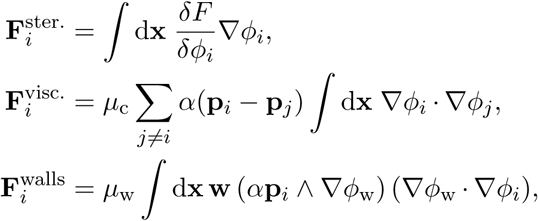

where **w** ≡ (−*∂_y_ϕ*_w_*,∂_x_ϕ*_w_) is a direction parallel to the wall, and *µ*_c_ and *µ*_w_ are dry-friction coefficients of the cells with other cells and the wall. Note that only cell-interfaces (where V*ϕ_i_* ≠ 0) contribute to the integrals. This form of the steric force is not unique but is well-motivated thermodynamically ^51,55^. Note that the steric force is normal to the interface while the viscous friction force is parallel to it. In practice, these two forces corresponds to two different cellular mechanisms: the gradient of tension at cell-cell contacts and the forces exerted on focal adhesions during frictional cell movements.

### Simulation details

We simulated equation (2) using a finite difference scheme on a square lattice with a predictor-corrector step. Throughout this article, we used the following numerical values for the simulation parameters: *λ* = *R* = 8, *γ* =0.05, *µ* =0.03, *κ* =0.4, *ξ* = 1, *α* =0.1. The phase-field representing the walls is set to have an exponentially falling profile *ϕ*_w_ = exp(−2*d*), where *d* is the distance to the closest wall and *κ*_w_ = 2. We simulated square domains of edge length *W* = 50, 75, 100, 125, 150 lattice sites with *N* = 15, 34, 60, 94, 135 cells and set *µ*_w_ = 0 with *D* =0.01, *J* =0.03, *µ*_c_ =0.05, and *µ*_w_ = 0. We simulated rectangular domains of length *L* = 600 and width *W* = 50, 75, 100, 125, 150 lattice sites with *N* = 180, 270, 360, 450, 540, 630, 720 cells. We introduced friction with the walls *µ*_w_ =0.2 and set *D* =0.2, *J* =0.5, and *µ*_c_ =0.5 in order to shorten the computation time. We have made sure that using these parameters in the square domains did not significantly affect our analysis. In particular, we also observe sustained oscillations in square confinement in this case.

## Supporting information

## Acknowledgements

GP, JdA, SB, RMM and BL gratefully acknowledge financial supports from the Human Frontier Science Programme (grant RGP0040/2012), the European Research Council under the European Union’s Seventh Framework Programme (FP7/2007-2013)/ERC grant agreement no 617233, the LABEX ‘Who am I?’, the Fondation ARC pour la Recherche contre le Cance and the French National Research Agency (ANR grant). RM is supported by grant P2EZP2 165261 of the Swiss National Science Foundation.

## References

[1] Friedl, P. & Gilmour, D. Collective cell migration in morphogenesis, regeneration and cancer. Nat. Rev. Mol. Cell Biol. 10, 445–457 (2009).

[2] Cetera, M., et al. Epithelial rotation promotes the global alignment of contractile actin bundles during *Drosophila* egg chamber elongation. Nat. Commun. 5, 5511 (2014).

[3] Park, S., et al. Tissue-scale coordination of cellular behaviour promotes epidermal wound repair in live mice. Nat. Cell Biol. 19, 155–163 (2017).

[4] Montell, D.J. Border cell migration: the race is on. Nat. Rev. Mol. Cell Biol. 4, 13–24 (2003).

[5] Martin, A.C., Gelbart, M., Fernandez-Gonzalez, R., Kaschube, M. & Wieschaus, E.F. Integration of contractile forces during tissue invagination. J. Cell Biol. 188 735–749 (2010).

[6] Roh-Johnson, M. et al. Triggering a Cell Shape Change by Exploiting Pre-Existing Actomyosin Contractions. Science 335, 1232–1235 (2012).

[7] Serra-Picamal, X. et al. Mechanical waves during tissue expansion. Nat. Phys. 8, 628–634 (2012).

[8] Tlili, S., et al. Waves in cell monolayer without proliferation: density determines cell velocity and wave celerity. *arXiv*:1610.05420 (2016).

[9] Londono C., et al. Nonautonomous contact guidance signaling during collective cell migration. Proc. Natl Acad. Sci. USA 111, 1807–1812 (2014).

[10] Reffay, M. et al. Interplay of RhoA and mechanical forces in collective cell migration driven by leader cells. Nat. Cell Biol. 16, 217–223 (2014).

[11] Woods, M., et al. Directional Collective cell migration emerges as a property of cell interactions. PLoS One 9, e104969 (2014).

[12] Sunyer, R. et al. Collective cell durotaxis emerges from long-range intercellular force transmission. Science 353, 1157–1161.

[13] Haigo, S.L. & Bilder, D. Global tissue revolutions in a morphogenetic movement controlling elongation. Science 331, 1071–1074 (2011).

[14] Friedl, P., Locker, J., Sahai, E. & Segall, J.E. Classifying collective cancer cell invasion. Nat. Cell Biol. 14, 777–783 (2012).

[15] Wolf, K. et al. Physical limits of cell migration: Control by ECM space and nuclear deformation and tuning by proteolysis and traction force. J. Cell Biol. 7, 1069–1084 (2013).

[16] Lecuit, Th., & Lenne, P.-F. Cell surface mechanics and the control of cell shape, tissue patterns and morphogenesis. Nat. Rev. Mol. Cell Biol. 8, 633–644 (2007).

[17] Vedula, S.R.K., Leong, M.C. et al. Emerging modes of collective cell migration induced by geometrical constraints. Proc. Natl. Acad. Sci. USA 109, 12974–12979 (2012).

[18] Szabo, A., et al. In vivo confinement promotes collective migration of neural crest cells. J. Cell. Biol. 213, 543–555 (2016).

[19] Heisenberg, C.P. & Bellaïche, Y. Forces in tissue morphogenesis and patterning. Cell 153, 948–962 (2013).

[20] Balaji, R., Bielmeier, Ch., et al. Calcium spikes, waves and oscillations in a large, patterned epithelial tissue. Sci. Rep. 7, 42786 (2017).

[21] He, L., Wang, X., Tang, H.L. & Montell, D.J. Tissue elongation requires oscillating contractions of a basal actomyosin network. Nat. Cell Biol. 12, 1133–1142 (2010).

[22] Hubaud, A., Regev, I., Mahadevan, L. & Pourquié, O. Excitable dynamics and Yap-dependent mechanical cues drive the segmentation clock. Cell 171, 668–682 (2017).

[23] Doxzen, K. et al. Guidance of collective cell migration by substrate geometry. Integr. Biol. 5, 1026–1035 (2013).

[24] Deforet, M., Hakim, V., Yevick, H.G., Duclos, G. & Silberzan, P. Emergence of collective modes and tri-dimensional structures from epithelial confinement. Nat. Commun. 5, 3747 (2014).

[25] Notbohm, J. et al. Cellular contraction and polarization drive collective cellular motion. Biophys. J. 110, 2729–2738 (2016).

[26] Dupont, S. et al. Role of YAP/TAZ in mechanotransduction. Nature 474, 179–183 (2011).

[27] Choquet, D., Felsenfeld, D.P. & Sheetz, M.P. Extracellular matrix rigidity causes strengthening of integrin-cytostkeleton linkages. Cell 88, 39–48 (1997).

[28] Gupta, M. et al. Adaptive rheology and ordering of cell cytoskeleton govern matrix rigidity sensing. Nat. Commun. 6, 7525 (2015).

[29] Aragona, M.C. et al. A mechanical checkpoint controls multicellular growth through YAP/TAZ regulation by actin-processing factors. Cell 154, 1047–1059 (2013).

[30] Moroishi, T., Hansen, C. G., & Guan, K. L. The emerging roles of YAP and TAZ in cancer. Nat. Rev. Cancer 15, 73–79 (2015).

[31] Angelini, Th. E. et al. Glass-like dynamics of collective cell migration. Proc. Natl. Acad. Sci. USA 108, 4714–4719 (2011).

[32] Garcia, S., et al. Physics of active jamming during collective cellular motion in a monolayer. Proc. Natl. Acad. Sci. USA 112, 15314–15319 (2015).

[33] Fütterer, C., Colombo, C., Jülicher, F & Ott, A. Morphogenetic oscillations during symmetry breaking of regenerating Hydra vulgaris cells. Europhys. Lett. 64, 137–143 (2003).

[34] Lauschke, V., Tsiairis, C.D., François, P. & Aulehla, A. Scaling of embryonic patterning based on phase-gradient encoding. Nature 493, 101–105 (2013).

[35] Shih, N.P., François, P., Delaune, E.A. & Amacher, S.L. Dynamics of the slowing segmentation clock reveal alternating two-segment periodicity. Development 142, 1785–1793 (2015).

[36] Mercker, M., Köthe, A. & Marciniak-Czochra, A. Mechanochemical symmetry breaking in *Hydra* aggregates. Biophys. J. 108, 2396–2407 (2015).

[37] Hillenbrand, P., Gerland, U. & Tkacik, G. Beyond the French flag model: exploiting spatial and gene regulatory interactions for positional information. PLoS ONE 11, e0163628 (2016).

[38] Beaupeux, M. & François, P. Positional information from oscillatory phase shifts: insights from in silico evolution. Phys. Biol. 13, 036009 (2016).

[39] Dembo M., & Wang, Y.-L. Stresses at the Cell-to-Substrate Interface during Locomotion of Fibroblasts. Biophys. J. 76, (4) 2307–2316 (1999).

[40] Trepat, X., et al. Physical forces during collective cell migration. Nat. Phys. 5, 426–430 (2009).

[41] Broussard, J.A. et al. The desmoplakin/intermediate filament linkage regulates cell mechanics. Mol. Biol. Cell 28, 3156–3164 (2017).

[42] Hatzfeld, M. & Magin, T.m. Desmosomes and intermediate filaments: their consequences for tissue mechanics. Cold Spring Harb. Perspect. Biol. 9, a029157 (2017).

[43] Yonemura, Sh., Wada, Y., Watanabe, T., Nagafuchi, A., & Shibata, M. *α*-Catenin as a tension transducer that induces adherens junction development. Nat. Cell. Biol. 12 (6), 533–542 (2010).

[44] Yao, M., Qui W. et al. Force-dependent conformational switch of *α*-catenin controls vinculin binding Nat. Commun. 5, 4525 (2014).

[45] Saw, Th. B., Doostmohammadi, A. et al. Topological defects in epithelia govern cell death and extrusion. Nature 544, 212–216 (2017).

[46] Nier, V. et al. Inference of Internal Stress in a Cell Monolayer. Biophys. J. 110, 1625–1635 (2016).

[47] Huveneers, S. & de Rooij, J. Mechanosensitive systems at the cadherin-F-actin interface. J. Cell Sci. 126, 403–413 (2013).

[48] Benham-Pyle, B. W., Pruitt, B. L., & Nelson, W. J. Mechanical strain induces E-cadherin-dependent Yap1 and *β*-catenin activation to drive cell cycle entry. Science 348, 1024–1027 (2015).

[49] Elosegui-Artola, A. et al. Force triggers YAP nuclear entry by regulating transport across nuclear pores. Cell 171, 1397–1410 (2017).

[50] Aranson I. S. Physical Models of Cell Motility. Springer International Publishing (2015)

[51] Palmieri, B., Bresler, Y., Wirtz, D., & Grant, M. Multiple scale model for cell migration in monolayers: Elastic mismatch between cells enhances motility. Sci. Rep. 5, 11745 (2015).

[52] Pilhwa Lee, Charles W. Wolgemuth. Crawling Cells Can Close Wounds without Purse Strings or Signaling. PLOS Comput. Biol. 7, e1002007 (2011).

[53] Szabó, B., Szöllösi, G. J., Gönci, B., Jurányi, Zs., Selmeczi, D., & Vicsek, T. Phase transition in the collective migration of tissue cells: Experiment and model. Phys. Rev. E 74, 061908 (2006).

[54] Henkes, S., Fily, Y., & Marchetti, M.C. Active jamming: Self-propelled soft particles at high density. Phys. Rev. E 84, 040301 (2011).

[55] Bitbol, A.-F., & Fournier, J.-B. Forces exerted by a correlated fluid on embedded inclusions Phys. Rev. E 83, 061107 (2011).

[56] Stramer, B., & Mayor, R. Mechanisms and *in vivo* functions of contact inhibition of locomotion. Nat. Rev. Mol. Cell Biol. 18, 43–55 (2017).

[57] Vedula, S.R.K., Peyret, G. et al. Epithelial bridges maintain tissue integrity during collective cell migration. Nat. Mater. 13, 87–96 (2014).

[58] Vedula, S.R.K., Peyret, G. et al. Mechanics of epithelial closure over non-adherent environments. Nat. Commun. 6, 6111 (2015).

[59] Hubaud, A. & Pourquié, O. Signalling dynamics in vertebrate segmentation. Nat. Rev. Mol. Cell Biol. 15, 709–721 (2014).

[60] Ruiz, S.A. & Chen, C.S. Emergence of patterned stem cell differentiation within multicellular structures. Stem Cells 26, 2921–2927 (2008).

[61] Zehnder, S.M., Wiatt, M.K., Uruena, J.M., Dunn, A.C. Sawyer, W.G. & Angelini, T.E. Multicellular density fluctuations in epithelial monolayers. Phys. Rev. E 92, 032729 (2015).

[62] Murakoshi, H., Lee, S.J. & Yasuda, R. Highly sensitive and quantitative FRET-FLIM imaging in single dendritic spines using improved non-radiative YFP. Brain Cell Biol. 36, 31–42 (2008).

[63] Basu, S., Totty, N.F., Irwin, M.S., Sudol, M. & Downward, J. Akt phosphorylates the Yes-associated protein, YAP, to induce interaction with 14-3-3 and attenuation of p73-mediated apoptosis. J. Mol Cell. 11, 11–23 (2003).

[64] Tsing, Q. ImageJ plugins for Traction Force Microscopy, https://sites.google.com/site/qingzongtseng/tfm, [Accessed: 2017-11-18].

[65] Malinverno, C. et al. Endocytic reawakening of motility in jammed epithelia. Nat. Mater. 16, 587–596 (2017).

