## Supplementary material for "Sustained oscillations of epithelial cell sheets"

### SI Appendix

#### SI Figures

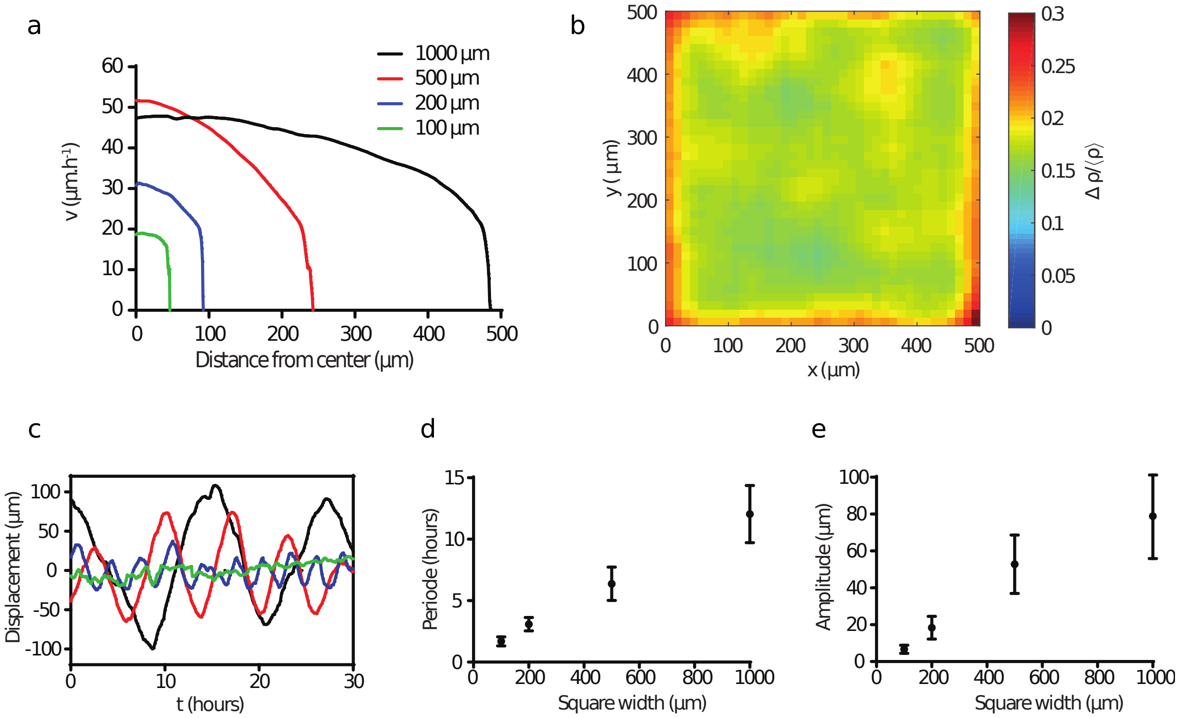


*Supplementary Figure 1:* **Oscillation dynamics**. **a**, *Average speed as a function of distance to the center for various square widths. The velocity drops sharply in a thin edge layer.* **b**, *Relative density fluctuations, measured with histone-GFP cells on* 500 µm *squares (average is taken over 20 squares). The most important deformations are concentrated on the edges as well.* **c**, *x-projection of single cell trajectories in squares of various sizes (same colours as in (a)).* **d** *and* **e**, *Oscillation properties of single cell trajectories: period and amplitude. Mean (SD) from* $n = 15, 15, 12$ *and* 9 *cells for* $W = 100, 200, 500$ *and* $1000$ µm *respectively from* $N \geqslant 3$ *experiments.*

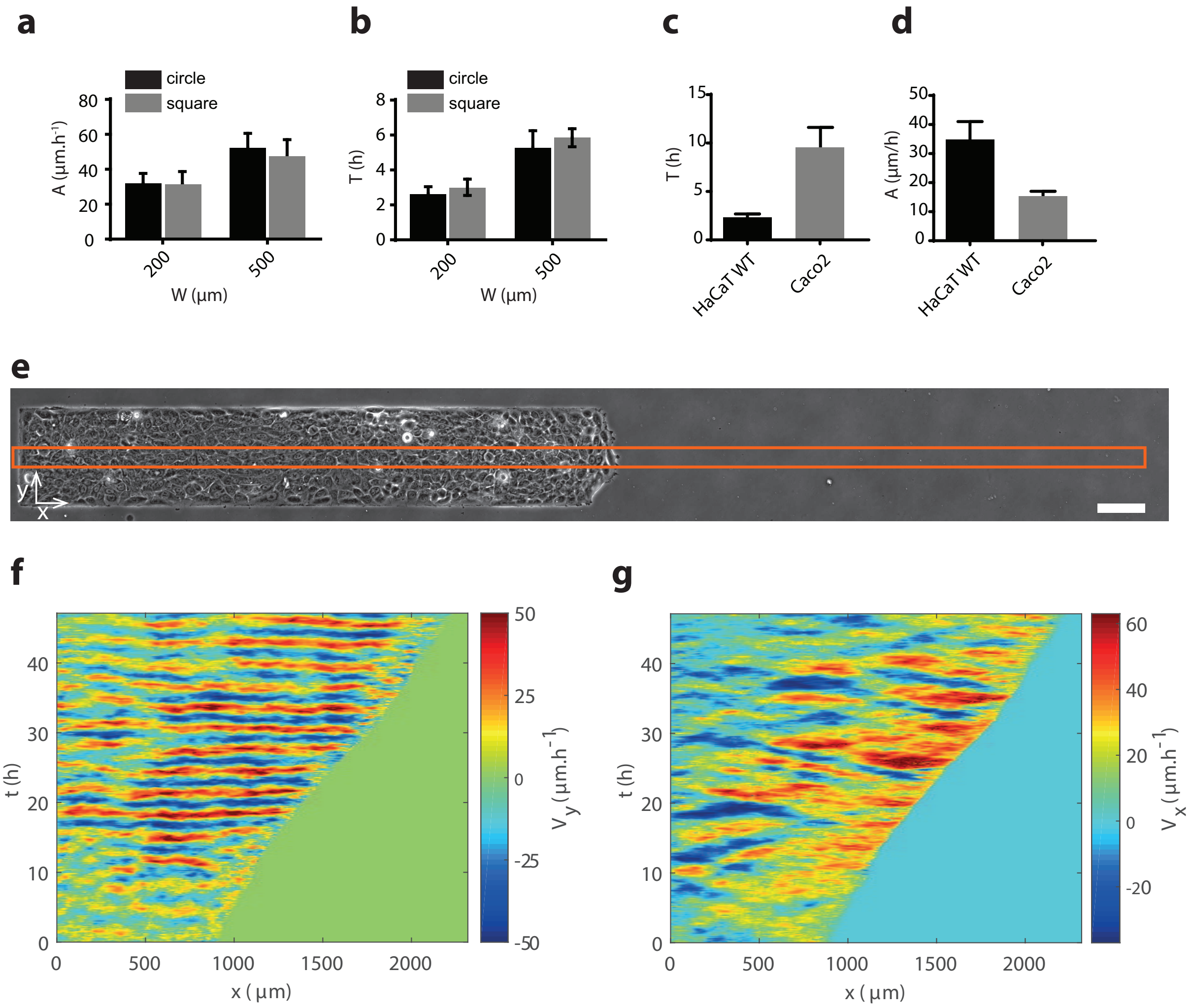


*Supplementary Figure 2:* **Coordinated oscillation is not a specific behaviour of HaCaT cells on squares**. **a, b**, *Comparison of oscillation amplitude $A$ and period $T$ for HaCaT cells on squares and disks of two different sizes.* **c, d**, *Comparison of oscillation amplitude $A$ and period $T$ for HaCaT and Caco2 cells on* 200 µm *squares.* **e**, *The cells were seeded in a* 200 µm*-wide band on a block of PDMS, and then allowed to migrate freely towards the right. The orange rectangle represents the ROI used to build the kymographs in* **f–g**. *Scale bar* 100 µm. **f**, *Kymograph of $V_y$, showing clear oscillations with strong coordination along the whole $x$-axis.* **g**, *Kymograph of $V_x$. The oscillations are centred around strictly positive value since the monolayer migrates to the right. n.s.: not significant, *$p < 0.05$, **$p < 0.01$, ***$p < 0.001$ from a two-sample Kolmogorov-Smirnov test.*

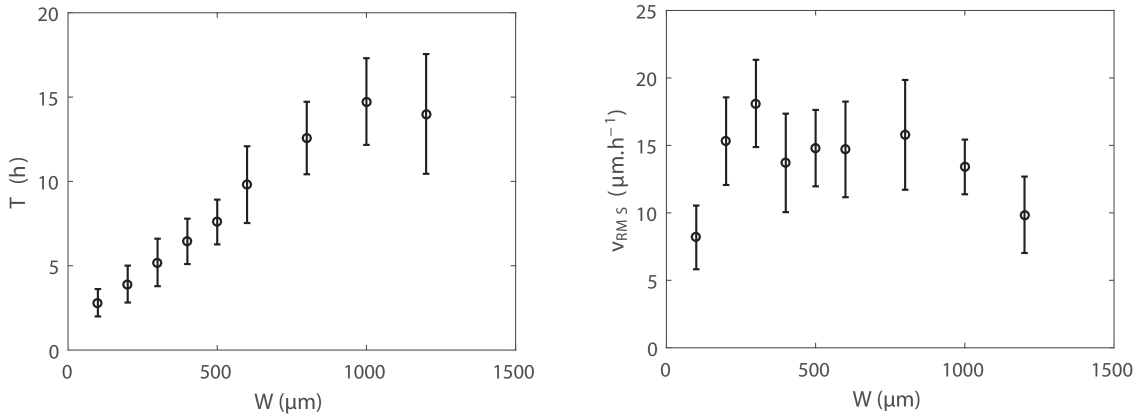


*Supplementary Figure 3:* **Size dependence of oscillations properties.** *Period $T$ oscillations and root-mean-square value of the collective velocity $v_{\mathrm{RMS}}$, as a function of the confinement size $W$. $n = 2761, 15, 10, 5, 16, 10, 6, 4$ squares for $W = 100, 200, 300, 400, 500, 600, 800, 1000, 1200\,\mu m$ respectively.*

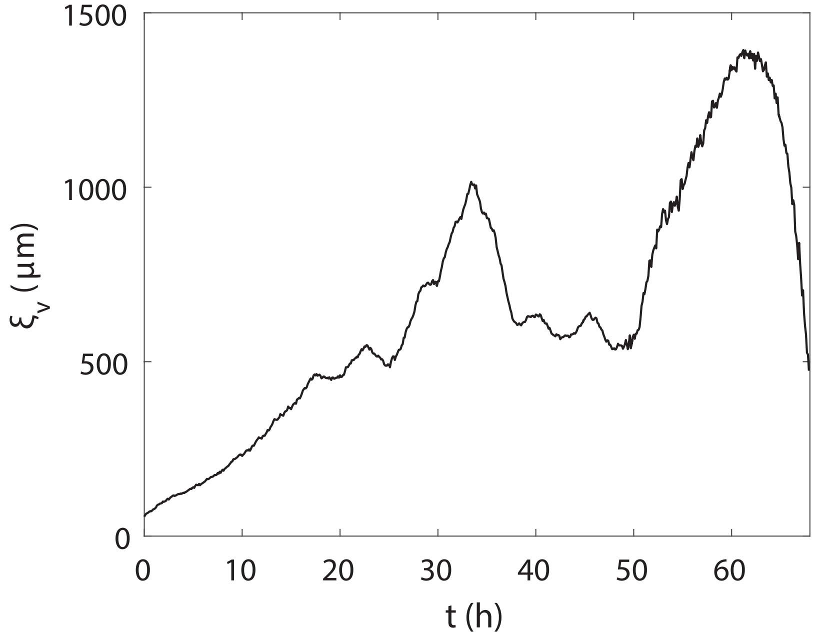


*Supplementary Figure 4:* **Movements at various time and length scales in unconstrained tissue**. *Correlation length versus time in an unconstrained confluent tissue of wild-type HaCaT cells. 90% and full confluence are reached at $t \simeq 20\,\mathrm{h}$ and $t \simeq 35\,\mathrm{h}$ respectively. Complex spatio-temporal patterns of collective motion emerge with typical values of $\xi_{\mathrm{v}}$ in the $600 - 1000\,\mu\mathrm{m}$ range.*

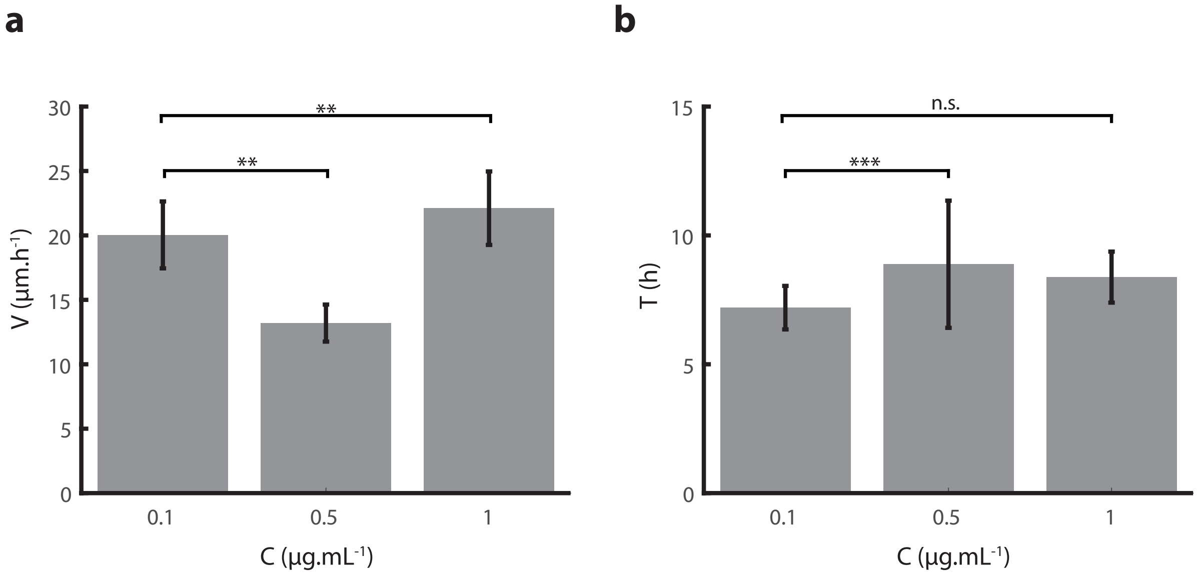


*Supplementary Figure 5:* **Verteporfin inhibition of YAP translationnal activity does not affect oscillations. a** *Mean speed* $V$ *and* **b** *period of oscillation of monolayers treated with various doses of verteporfin. n.s.: not significant,* $*p < 0.05$, $**p < 0.01$, $***p < 0.001$ *from a two-sample Kolmogorov-Smirnov test.*

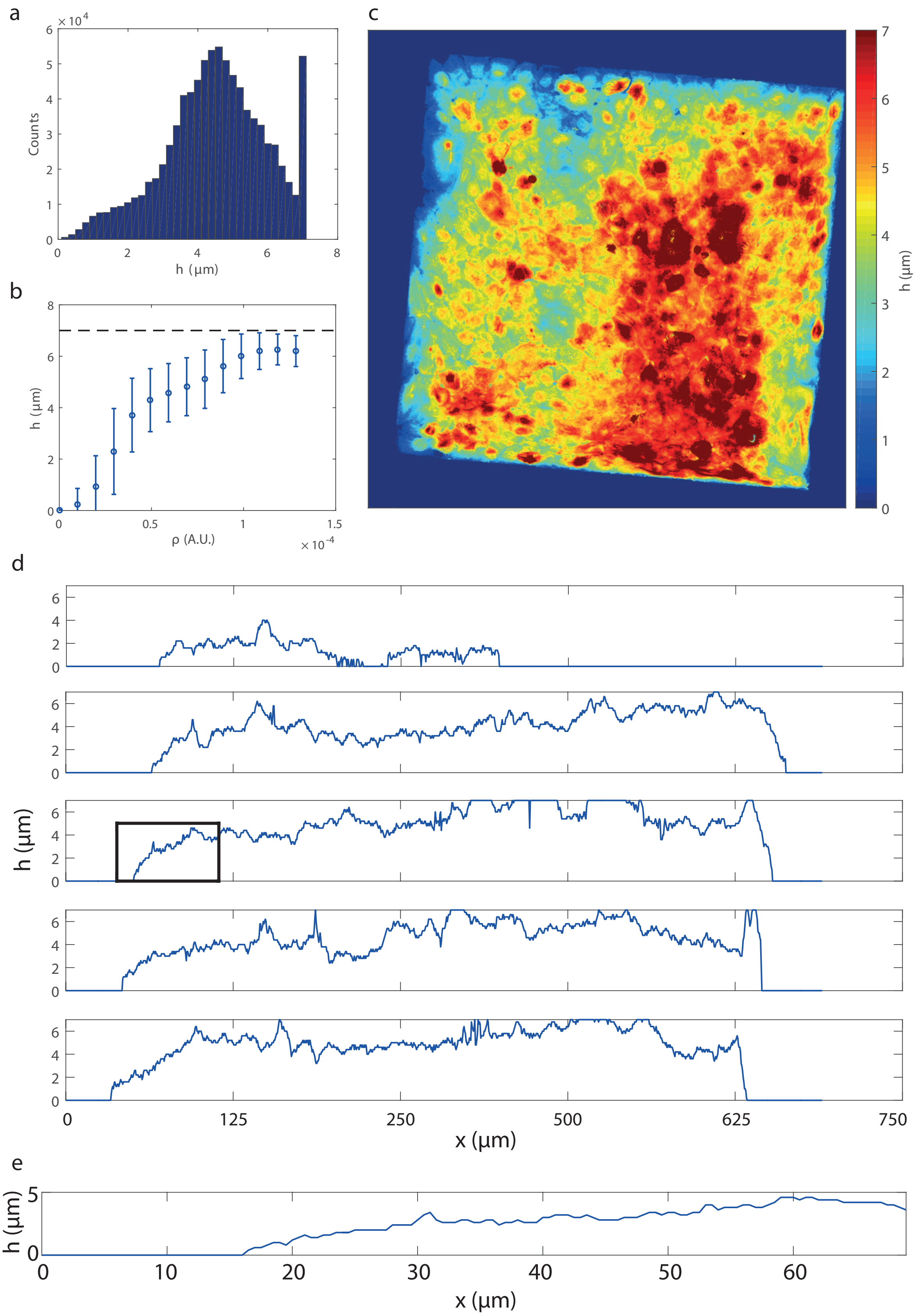


*Supplementary Figure 6:* **Monolayer height.** **a** *Distribution of height on a monolayer confined on a 500* μm *square.* **b** *Height as a function of local cell density (same data as in Figure 3).* **c** *Height map, measured from confocal z-stack of immunolabeled monolayer of HaCaT cells confined on a 500* μm *square (same data as in Figure 3).* **d** *Profiles of height at various y.* **e** *Close-up on the edge of the monolayer (black box on the third profile) showing h and x on the same scale.*

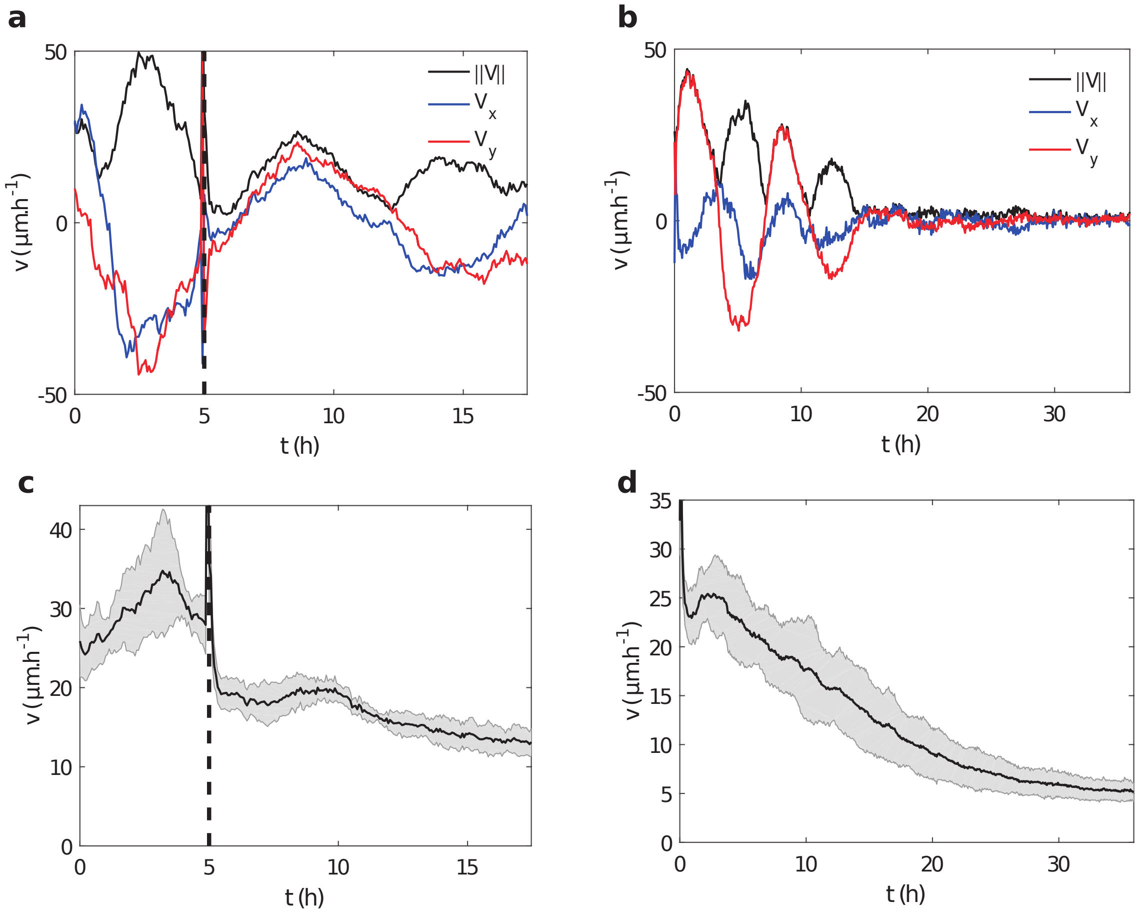


*Supplementary Figure 7:* **Inhibition of cytoskeleton activity affects the motion but not directly the oscillations**. ***a***, *Example oscillation of HaCaT cells in a 500* µm *square. Blebbistatin is added at* $t = 5\,h$ *(dashed vertical line).* ***b***, *Example oscillation of HaCaT cells in a 500* µm *square. CK666 is added at* $t = 0\,\mathrm{h}$. ***c***, *Average speed of HaCaT cells in 500* µm *squares. Blebbistatin is added at* $t = 5\mathrm{h}$ *(dashed vertical line). Mean (SD) of* $n = 5$ *squares.* ***d***, *Average speed of HaCaT cells in 500* µm *squares. CK666 is added at* $t = 0\mathrm{h}$. *Mean (SD) of* $n = 15$ *squares.*

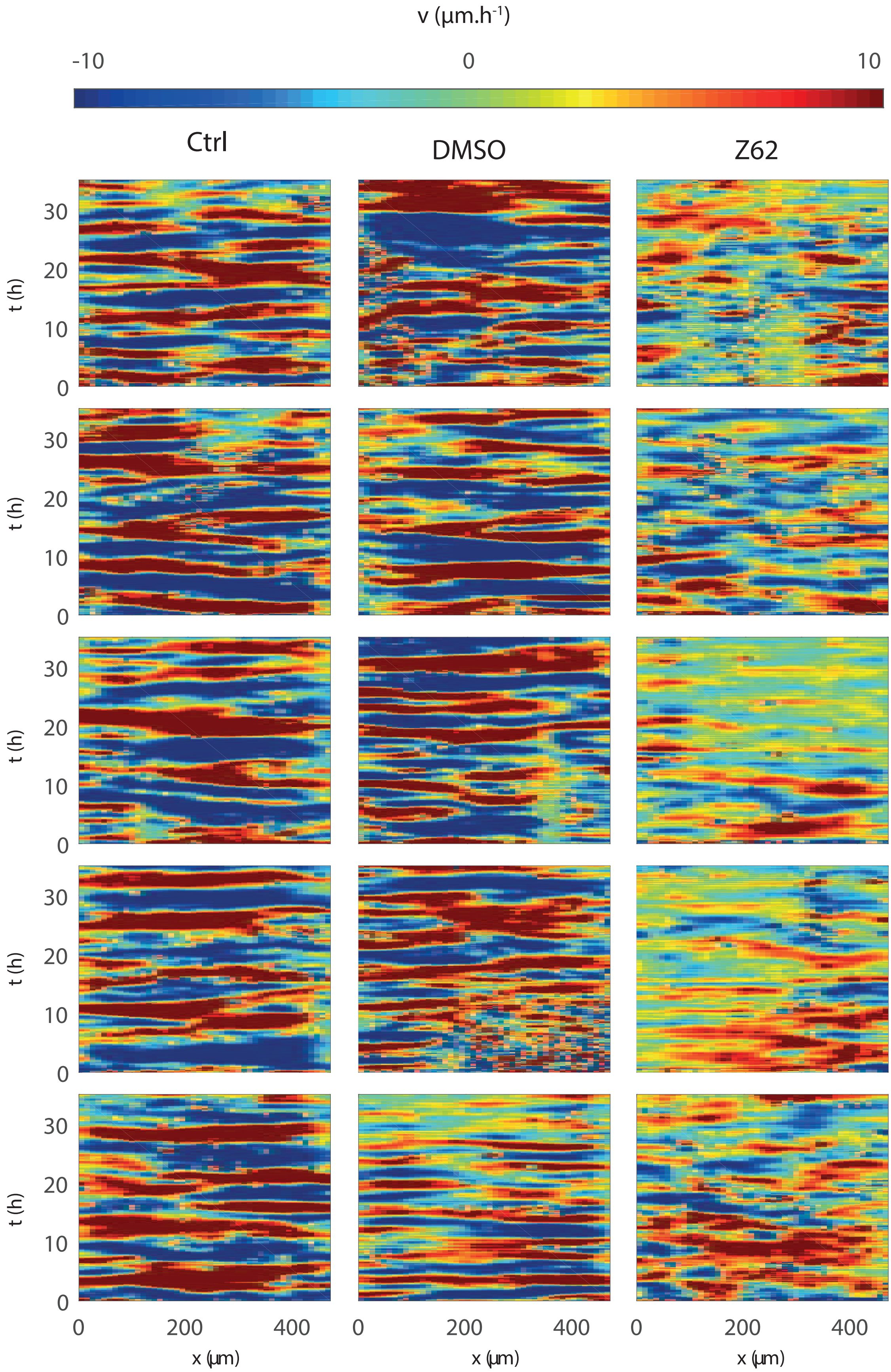


*Supplementary Figure 8:* **Rac1 inhibition suppresses oscillations.** *Typical kymographs of $v_x$ for HaCaT cells in control condition, treated with DMSO (control) and Z62954982 Rac1 inhibitor. Not only does the overall cell speed reduce, but the movement becomes less coordinated and oscillations disappear.*

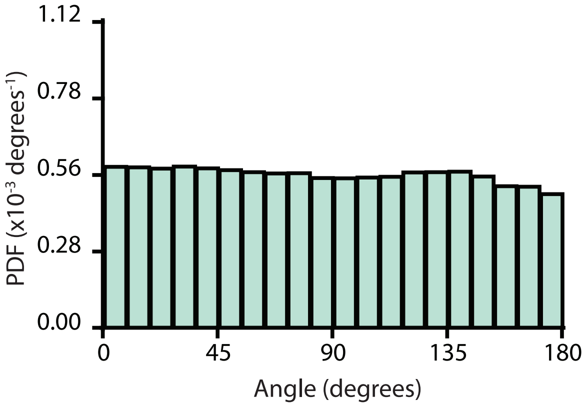


*Supplementary Figure 9:* **Histogram of the angle between traction force and velocity.**

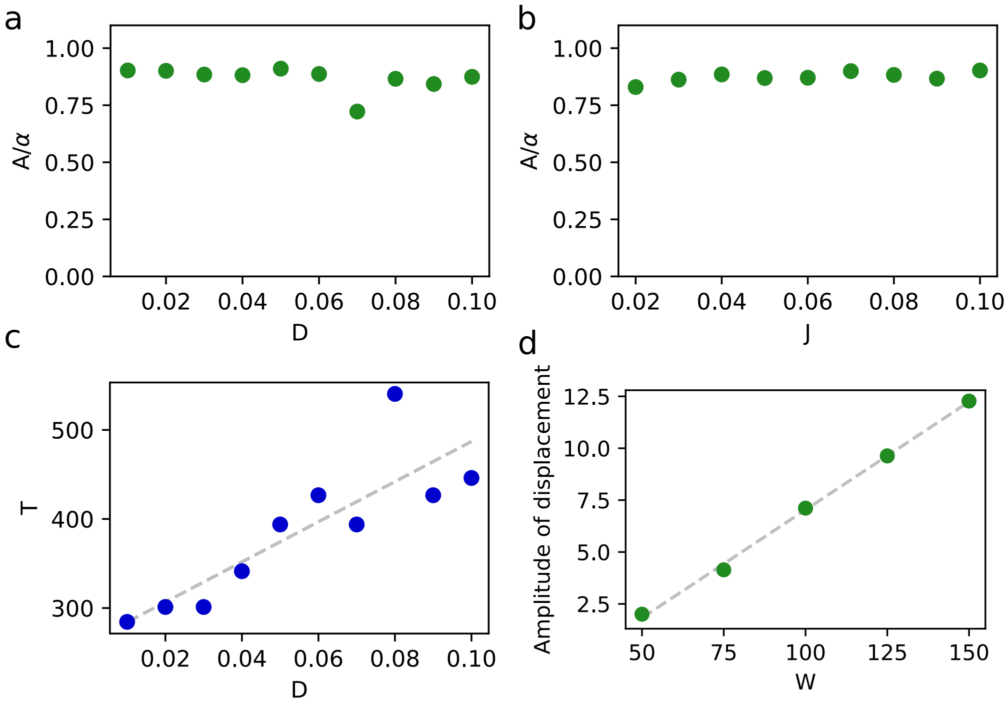


*Supplementary Figure 10:* **Dependence of oscillation properties on model parameters.** **a** *and* **b**, *Dependence of the amplitude of the oscillations on the alignment parameters of equation (1). The amplitude is not affected by changes in $D$ and $J$.* **c**, *Dependence of the period of oscillation on the noise parameter $D$. Grey dashed line is a least-square fit of $T \propto D$.* **d**, *Amplitude of displacement, measured as the amplitude of oscillation of the centroid of the individual cells centres.*

#### SI Videos

Supplementary Video 1:
SupplementaryVideo-1.mp4
Layer-scale coordinated movements of HaCaT cells in square confinement. A few cells have been tracked manually. Scale bar 100 μm.

Supplementary Video 2:
SupplementaryVideo-2.mp4
Layer-scale coordinated movements of HaCaT cells in rectangular confinement. Scale bar 100 μm.

Supplementary Video 3:
SupplementaryVideo-3.mp4
YAP-GFP HaCaT cell moving with changes of cell size and related relocalisation of YAP. Scale bar 10 μm.

Supplementary Video 4:
SupplementaryVideo-4.mp4
Uncoordinated movements of $\alpha$-catenin KD HaCaT cells in square confinement. A few cells have been tracked manually. Scale bar 100 μm.

Supplementary Video 5:
SupplementaryVideo-5.mp4
Simulations of confluent monolayers in square confinements show the emergence of sustained oscillations

Supplementary Video 6:
SupplementaryVideo-6.mp4
Closure of a non-adherent gap by HaCaT cells. The epithelium exhibits coordinated oscillations on a large scale, with a strong impact on the closure dynamics.
